# Myeloid Tribbles 1 induces early atherosclerosis via enhanced foam cell expansion

**DOI:** 10.1101/615872

**Authors:** Jessica M Johnston, Adrienn Angyal, Robert C Bauer, Stephen Hamby, S Kim Suvarna, Kajus Baidžajevas, Zoltan Hegedus, T Neil Dear, Martin Turner, The Cardiogenics Consortium, Heather L Wilson, Alison H Goodall, Daniel J Rader, Carol C Shoulders, Sheila E Francis, Endre Kiss-Toth

**Author notes:** Membership of the Cardiogenics Consortium appears in the Supplementary Materials.

## Abstract

Macrophages drive atherosclerotic plaque progression and rupture, hence attenuating their atherosclerosis-inducing properties holds promise for reducing coronary heart disease (CHD). Recent studies in mouse models have demonstrated that Tribbles 1 (Trib1) regulates macrophage phenotype and shows that *Trib1* deficiency increases plasma cholesterol and triglyceride levels, suggesting that reduced *TRIB1* expression mediates the strong genetic association between the *TRIB1* locus and increased CHD risk in man. However, we report here that myeloid-specific *Trib1* (m*Trib1*) deficiency reduces early atheroma formation and that m*Trib1* transgene expression increases atherogenesis. Mechanistically, m*Trib1* increased macrophage lipid accumulation and the expression of a critical receptor (OLR1), promoting oxidized low density lipoprotein uptake and the formation of lipid-laden foam cells. As *TRIB1* and *OLR1* RNA levels were also strongly correlated in human macrophages, we suggest that a conserved, TRIB1-mediated mechanism drives foam cell formation in atherosclerotic plaque and that inhibiting mTRIB1 could be used therapeutically to reduce CHD.

## Introduction

Atherosclerosis, a progressive disease of arterial blood vessels and the main underlying cause of stroke, myocardial infarction and cardiac death (1), is initiated by the conversion of plaque-macrophages to cholesterol-laden foam cells (2) in the arterial intima (3). In the early-stage atherosclerotic plaque this transformation is induced by the uptake of both LDL-C and oxidized (ox)LDL (2, 4), which may serve a beneficial purpose (3); but unrestrained, the crucial function of plaque-macrophages in resolving local inflammation is compromised and the development of unstable, advanced lesions ensues (3). Importantly, it has been shown that foamy macrophages are not only less effective in clearing apoptotic cells (5) they are also more prone to apoptosis (6), thus increasing secondary necrosis and the release of cellular components and lipids that ultimately form the necrotic core of advanced plaques. As such, there have been investigations into the identities of macrophage-specific proteins that induce lipid accumulation. Thus, myeloid-lipoprotein lipase (LPL), for example, has been shown to enhance the retention of LDL-C and triglyceride-rich remnant particles within the artery wall (7) and induce foam cell formation (8); while the scavenger receptor, oxidized low-density lipoprotein receptor 1 (OLR1) has been found to internalize oxLDL (9), promoting not only lipid accumulation and growth but also the survival of macrophage-foam cells (10). Conversely, myeloid-*ApoE* expression has been shown to promote HDL-mediated cholesterol efflux (11) and macrophage switching from a pro-inflammatory (M1) to an alternatively (M2) activated phenotype (12). However, significant advances in the development of CVD therapeutics await the identification of an apical regulator(s) that acts in a coordinated manner on the multiple downstream processes governing lipid accumulation, as well as atherogenicity of plaque-resident macrophages.

Tribbles-1 has been detected in murine plaque-resident macrophages (13) and variants at the *TRIB1* locus have been associated with increased risk of hyperlipidemia and atherosclerotic disease in multiple populations (14–16). However, no study had examined the macrophage-specific cellular processes dependent on m*Trib1* expression and how these tally with the assumed athero-protective properties of this pseudokinase. At the whole-body level, one study has shown that *Trib1* deficient mice have markedly reduced numbers of M2-like (F4/80^+^ MR^+^) macrophages in multiple organs, including adipose tissue (17). As such, these studies strongly implicated that loss of macrophage-*Trib1* expression within the arterial wall would lead to excessive atherosclerotic plaque inflammation and/or impair inflammation resolution and promote atheroma formation. Moreover, in hepatocytes *Trib1* suppresses VLDL production and *de novo* lipogenesis (16), indicating that the association between variants at the *TRIB1* locus and atherosclerotic disease (14–16) relates to loss of TRIB1 activity.

In the current study, we found that contrary to expectations, myeloid-specific knockout of *Trib1* is athero-protective, while m*Trib1* expression is detrimental. In brief, *Trib1* increased OLR1 RNA and protein expression, ox-LDL uptake, foamy macrophage formation and atherosclerotic burden in two distinct mouse models of human disease. The expression of these two genes, as well as those of *LPL* and *SCARB1* (which mediates selective HDL-cholesterol uptake (18)), are also tightly linked in human macrophages. Collectively, our studies reveal an unexpected beneficial effect for selectively silencing *Trib1* in arterial plaque macrophages.

## Results

### Myeloid-Trib1 Increases Atherosclerosis Burden

Immunostaining of a human coronary atheroma detected Tribbles 1 in the arterial wall, including in macrophages (**Fig 1A**). We therefore examined the impact of macrophage Tribbles 1 expression on atherogenesis by creating mice expressing low, WT and elevated levels of myeloid *Trib1* as outlined in **Fig 1 B – F**. Although previous studies have demonstrated that global *Trib*1 KO significantly increases perinatal lethality (17) *Trib1*-floxed mice and myeloid-specific Trib1 knock-out (*Trib1*^mKO^) mice were fully viable and bred normally (**Fig S1 A-D**). Myeloid-specific *Trib1* transgenic (overexpressing, *Trib1*^mTg^) mice were also fully viable and bred normally. m*Trib1* RNA levels were substantially lower in *Trib1*^mKO^ than in floxed WT littermates, as judged by RT-qPCR assays performed on bone marrow-derived macrophages (BMDMs) prepared from these animals (**Fig 1G**). As judged by eGFP expression, the bi-cistronic *Trib1* transgene was expressed in 78.43 ± 2.33% and 65.58 ± 0.92% of blood monocytes and peritoneal macrophages, respectively (**Fig 1G**) and overall, the transgene increased BMDM *Trib1* RNA levels by 2.49 ± 0.43 (SEM) fold (**Fig 1G**, fourth panel). Consistent with previous findings (19) the transgene was also expressed in neutrophils, which form a minor component of the immune cell population within very early-stage atherosclerotic lesions (20). Thus, we detected eGFP in 53.88 ± 2.41% and 34.93 ± 2.96% of *Trib1*^mTg^ blood and bone marrow CD11b^+^/Ly6C^-^/Ly6G^+^ cells, respectively compared to 25.95 ± 3.16% and 12.42 ± 2.01% in their CD11b^+^/Ly6C^+^/Ly6G^-^ monocyte counterparts (**Fig. S1 E - I)**. However, in marked contrast to the reported full-body *Trib1* knockout mouse (17), *Trib1*^mKO^ mice were not afflicted by reduced numbers of total, or individual, white blood cells (**Fig. 1H**) or by reduced macrophage numbers in their adipose tissue (F4/80^+^, **Fig. S2A**), liver (F4/80^+^, **Fig. S2B**) or spleen (F4/80^+^ and CD206^+^, **Fig. S2C**). Similar to *Trib1*^mKO^ mice, *Trib1*^mTg^ mice displayed no gross abnormalities and had WT numbers of white blood cells (**Fig. 1H).** Additionally, the sizes of their adipocytes (**Fig. S2A)**, liver (**Fig. S2B)** and splenic (**Fig. S2C**) macrophage populations were unaltered.

**Fig. 1.**
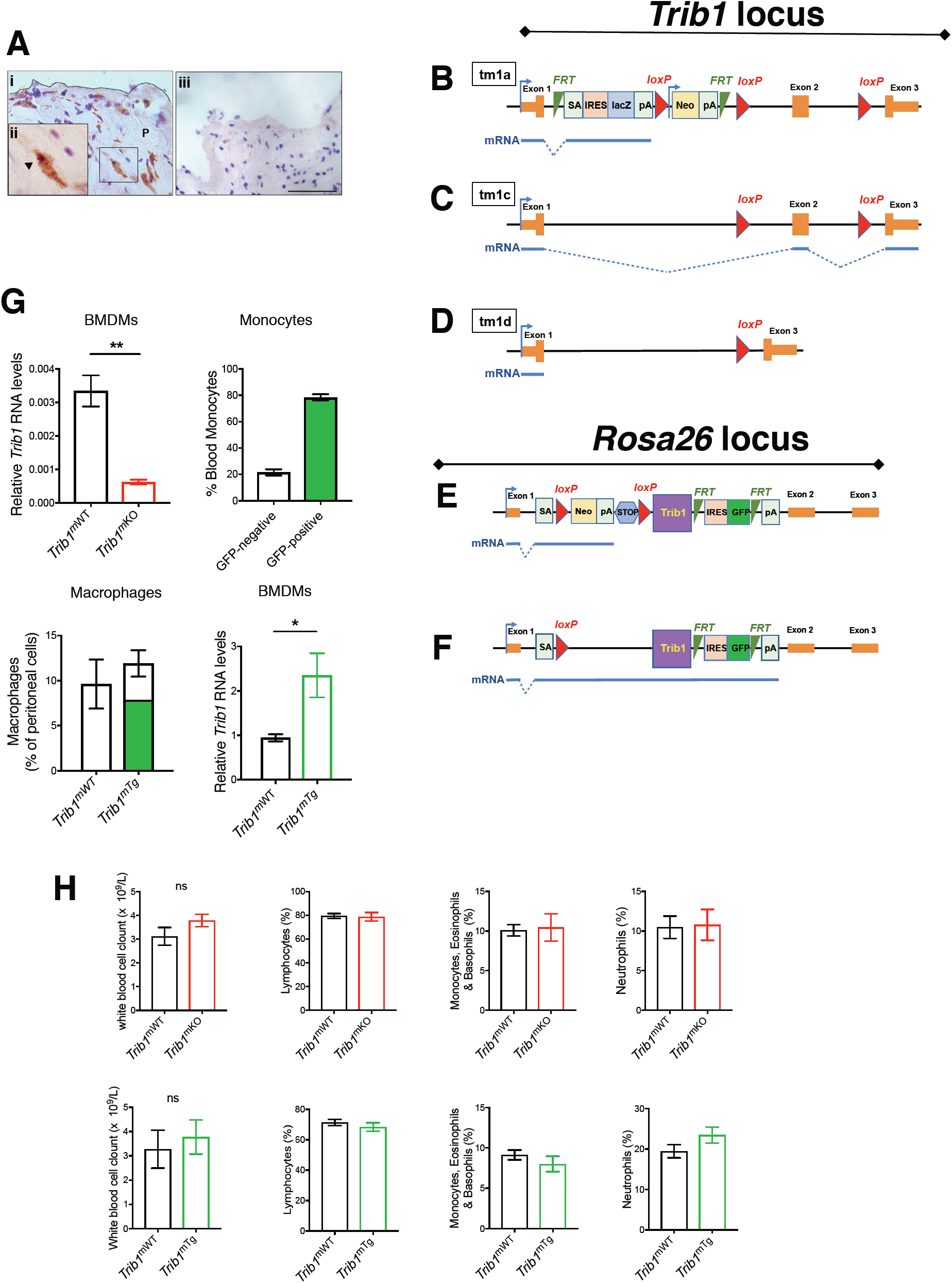
Generation and Characterization of Myeloid-specific Strains of *Trib1*-Knock-out and Transgenic Mice. ***(A*)** Immunohistochemistry of human atherosclerotic plaque (P). TRIB1 (red). CD68+ macrophages (brown). (**ii**) Magnification (x40) of boxed area highlighting one of the double positive cells. (**iii**) Isotype control (Scale: 50µm). **(*B*)** Schematic of the *Trib1* ‘knockout-first’ targeting construct used to produce null, conditional-ready/floxed (tm1c) and conditional-null (tm1d) alleles. Predicted transcripts below. FRT, flippase recognition target; SA, splice acceptor sequence; pA, polyadenylation signal; IRES, internal ribosomal entry site; LacZ, β-galactosidase; Neo, neomycin resistance gene. **(*C*)** The tm1c (*Trib1*^mWT^) allele created via flippase recombinase-mediated removal of the ‘gene-trap’ cassette. **(*D*)** The null/conditional-*Trib1* allele (tm1d) produced by crossing tm1c and Cre-expressing mice. Viability data for mice carrying these three *Trib1* alleles are provided in **Fig S1. (*E*)** *Rosa26*-STOP-Trib1-eGFP transgene construct used to produce *Trib1*^mWT^ and *Trib1*^mTg^ mice. **(*F*)** Cre-mediated excision of the STOP cassette enables transcription of the bi-cistronic *Trib1*-*eGFP* transcript from the endogenous *Rosa26* promoter (indicated by bent arrow). **(*G*)** *Trib1* RNA (relative to *Actb*) in bone marrow derived macrophages (BMDMs) from homozygous tm1c (i.e. *Trib1*^mWT^) and *Trib1*^*m*KO^ mice (n=3 per group). Percentages of monocytes and peritoneal macrophages from *Trib1*^mTg^ mice (n=3 per group) expressing eGFP. *Trib1* RNA levels in BMDMs from *Trib1*^mTg^ mice, expressed relative to *Trib1*^mWT^ (n=5-7 per group)**. (*H*)** Blood cell counts of mixed-gender *Trib1*^mKO^ (top panels) and *Trib1*^mTg^ (bottom panels) and their respective WT littermates (N= 5-6 per group). Data are mean ± SEM. Significance was determined by Student’s t-test, \**P* <0.05 and \*\**P* <0.01.

To address the contribution of myeloid *Trib1* in early atherosclerosis, we first transplanted bone marrow cells from the *Trib1*^mKO^ and *Trib1*^mTg^ mice and their respective controls (i.e. non-CRE, floxed KO and Tg alleles) into 12-13 weeks old lethally-irradiated male *ApoE*^-/-^ mice (**Fig 2A**). Thus, all recipient mice received *ApoE*^*+/+*^-bone marrow cells to mitigate the previously described effects of total ablation of this apolipoprotein on both classical/proinflammatory (M1) and alternative/anti-inflammatory (M2) polarization (12) and, to provide them with a physiologically important source of ApoE to aid normalisation of plasma cholesterol levels and of the lipoprotein profile in this otherwise extreme hyperlipidaemic mouse model of human atherosclerosis (12, 21). Following a seven-week recovery period, the chimeric mice were fed a Western diet containing 0.2% cholesterol for 12 weeks. At sacrifice, and consistent with expectations of the study design, the chimeric mice had relatively low plasma cholesterol levels for a mouse model of human atherosclerosis (**Fig S3A**).

**Fig. 2.**
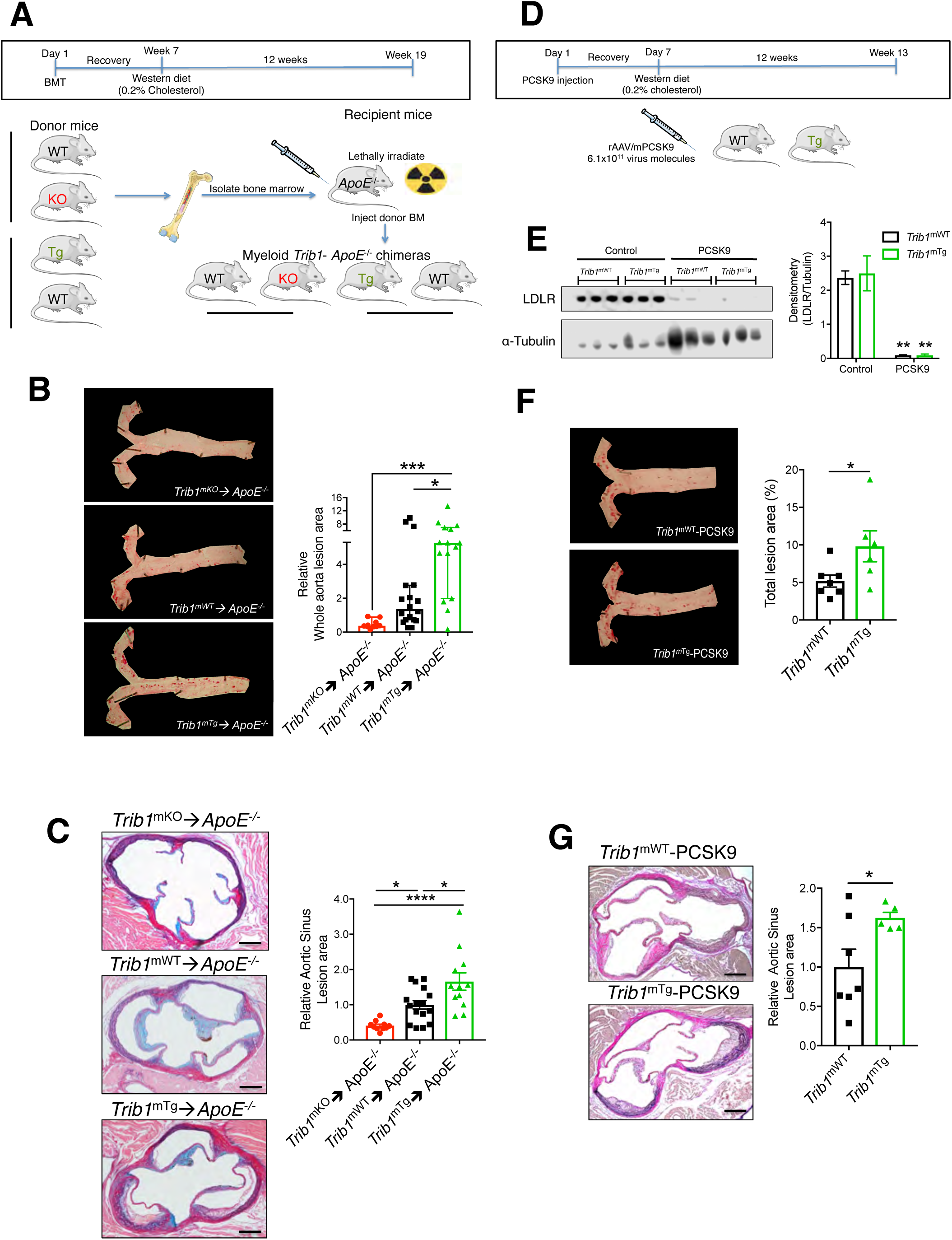
Myeloid *Trib1* Transgenic (*Trib1*^mTg^) Expression Increases Atherosclerosis Burden in Two Murine Models of Human Atherosclerosis. **(*A***) Schematic of the bone marrow transplant experiment. Bone marrow cells from myeloid-specific *Trib1* knock out (KO) and transgenic (Tg) mice and their respective wild-type (WT) controls were transplanted into *ApoE*^-/-^ recipients. **(*B*)** Representative *en face* Oil red O (ORO) staining of thoracic aortas from specified chimeras following the Western diet, and quantification. Lesion areas were calculated as percentages of the total surface area of the whole aorta and normalised (median, ± 95% CI) to *Trib1*^mWT^; n=10-18 per group. **(*C*)** Representative images of Elastic van Gieson-stained aortic sinus lesions and quantification (n=10-16 mice per group). **(*D*)** Strategy used to determine the consequences of m*Trib* transgene expression on atheroma burden in mice expressing adenovirus-produced proprotein convertase subtilisin/kexin 9 (PCSK9). **(*E*)** LDLR protein in cell lysates of liver samples harvested from specified mice was detected by Western Blot and quantified (n=3, per group). **(*F*)** Representative *en face* ORO staining of thoracic aortas from *Trib1*^mWT^-PCSK9 (top) and *Trib1*^mTg^-PCSK9 (bottom) mice. Lesion areas were calculated as percentages of the total surface areas of the whole aorta (n=6-7 per group). **(*G*)** Representative images of Elastic van Gieson-stained aortic sinus lesions of specified mice and quantification (n=5-7 per group). In **C** and **G**, scale bar = 200µm In ***B, C, F*** and ***G*** data are expressed relative to mWT. Data are mean ± SEM. Significance was determined by one-way ANOVA (***B, C***), two-way ANOVA (**e**) or student’s t-test (***F, G***). \**P* <0.05, \*\*\**P* <0.001, \*\*\*\**P* <0.0001.

Unexpectedly, we found less atherosclerosis in the thoracic aorta of *Trib1*^mKO^**→***ApoE*^-/-^ chimeras than in the control WT mice (**Fig. 2B, Fig. S4A).** Conversely, there was a significantly higher atheroma burden in the *Trib1*^mTg^**→***ApoE*^-/-^ mice (**Fig. 2B, Fig S4A**). Similarly, the lesions in the aortic sinus were on average smaller in the *Trib1*^mKO^**→***ApoE*^-/-^ mice and larger in the *Trib1*^mTg^**→***ApoE*^-/-^ mice (**Fig. 2C, Fig. S4B**). However, the collagen contents of the *Trib1*^mKO^**→***ApoE*^-/-^ and *Trib1*^mTg^**→***ApoE*^-/-^ in these “early-stage” plaques and their clinical pathology were comparable to those of the chimeric *Trib1*^mWT^**→***ApoE*^-/-^ mice (**Fig. S4C**). In short, we found that *mTrib1* expression increased the atherosclerotic burden of *ApoE*^-/-^ mice, despite having little impact on plasma LDL-cholesterol levels (**Fig S3A**).

To confirm that *mTrib1* accelerates the development of atherosclerosis (**Fig. 2B, C**) we created an LDL-receptor (*Ldlr*) knock-down model of human atherosclerosis (22) to induce hyperlipidaemia and atherosclerosis in otherwise WT mice (22). Specifically, mice were injected with an adeno-associated virus (rAAV8) encoding for proprotein convertase subtilisin/kexin 9 (**Fig. 2D**), which previous studies have shown lowers both hepatic and extrahepatic surface cell expression of the *Ldlr* (23). Following feeding a Western Diet for 12 weeks, this intervention produced comparable, highly significant reductions in LDLR protein levels in the *Trib1*^mTg^ and *Trib1*^mWT^ mice (**Fig. 2E**) and a similar degree of hyperlipidaemia (**Fig. S3B**). However, despite this and consistent with m*Trib1* expression increasing atheroma formation in *ApoE*^*-/-*^ mice, *Trib1*^mTg^ injected with rAAV8-Pcsk9 developed a significantly higher atherosclerotic burden in their aorta and aortic sinus than their similarly injected *Trib1*^mWT^ mice (**Fig. 2F-G**)

### Myeloid-Trib1 Increases Macrophage/Foam Cell Size in the Atherosclerotic Plaque

Next, we investigated the macrophage content and phenotype in the atherosclerotic plaque in each mouse models. This revealed that the aortic sinus lesions of *Trib1*^mKO^**→***ApoE*^-/-^ mice contained a much smaller MAC3^+^ immuno-reactive area than the chimeric *Trib1*^mWT^**→***ApoE*^-/-^ mice, while on average the *Trib1*^mTg^**→***ApoE*^-/-^ atheromas contained a marginally larger stained area (**Fig. 3A, B**). However, there was no preferential loss of YM1^+^ macrophages in the *Trib1*^mKO^**→***ApoE*^-/-^lesions (**Fig. 3B**), consistent with the finding that M2 polarization of *Trib1*-deficient BMDMs isolated from whole-body *Trib1*^mKO^ mice are compromised to a similar extent as M1 polarization (24). Additionally, we could not attribute the pro-atherogenic activity of myeloid *Trib1* expression to a preferential increase in the pro-inflammatory macrophage (NOS2^+^) content of *Trib1*^mTg^**→***ApoE*^-/-^ plaque (**Fig. 3B, panel 3**). Rather, the increased atherosclerosis in the *Trib1*^mTg^**→***ApoE*^-/-^ chimeras was attributable to a doubling of foam cell numbers (cells with characteristic foamy appearance and MAC3+ (**Fig S4D**)), and on average, these cells were also larger (**Fig. 3A, C, Fig S4D**). Likewise, there was no difference in the sizes of the macrophage populations in the *Trib1*^mTg^ -*Pcsk9* and *Trib1*^mWT^-*Pcsk9* mice atheromas (**Fig. 3D**) and despite the increased atherosclerotic burden in the transgenic animals **(Fig. 2F, G**), the atherosclerotic lesions of the *Pcsk9*-*Trib1*^mTg^ mice contained larger foam cells (**Fig. 3D**).

**Fig. 3.**
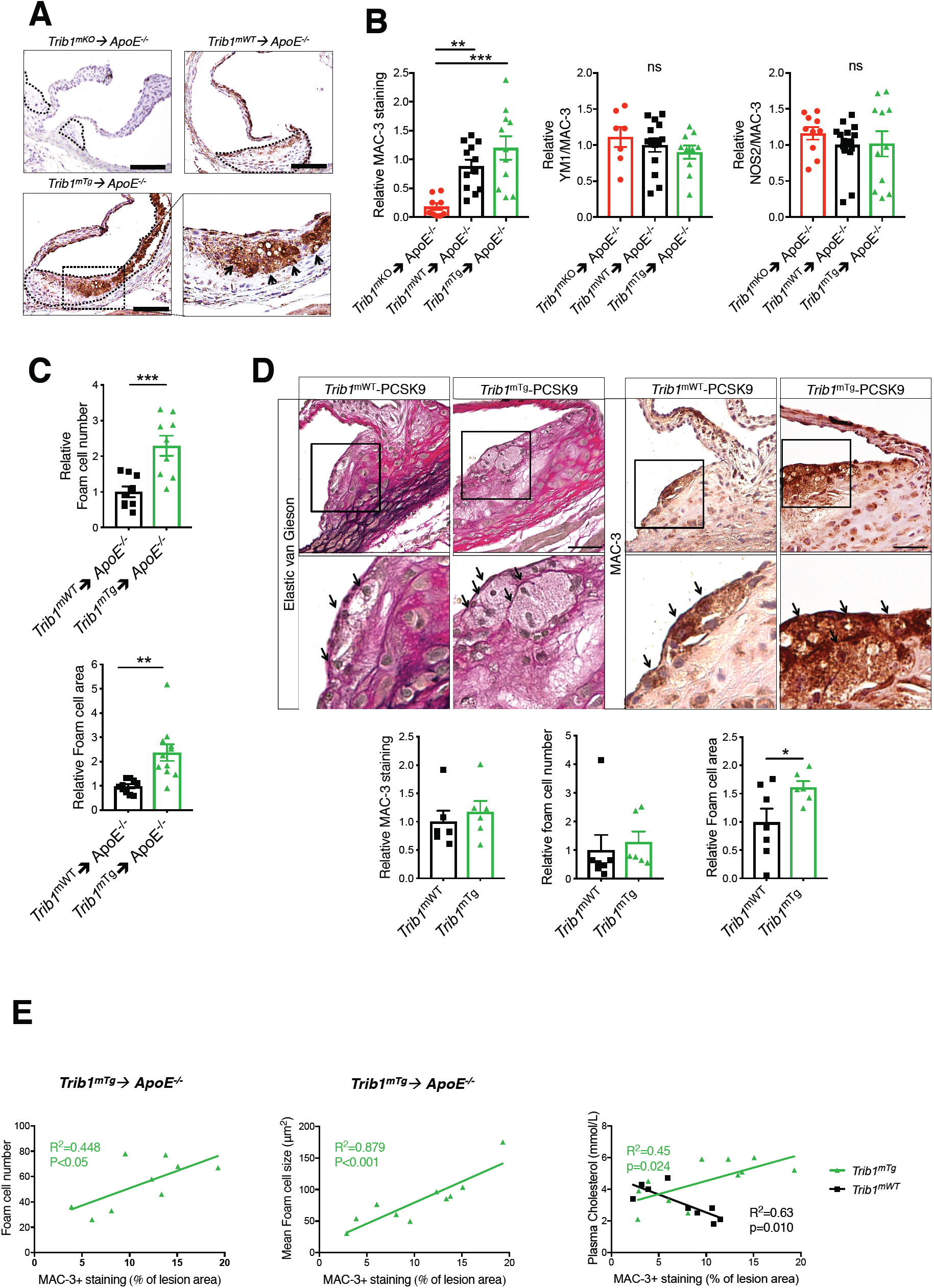
Myeloid-Trib1 Induces Foam Cell Expansion. *(A)* MAC-3 staining (brown) of representative cross-sections of the aortic sinus from specified mice (magnification x20, scale: 100µm). Dashed line indicates boundaries of lesions. Higher magnification (x40) of boxed area highlights the high number of foam cells (arrows) in the aortic sinus plaque of *Trib1*^***mTg***^**→***ApoE*^*-/-*^ mice. **(*B*)** Staining of aortic sinus lesions from specified chimeric mice with specified antibodies: First panel, MAC-3 (n= 9-12 per group). Second panel, YM1/MAC-3 double-positive cells (n= 7-14 per group). Third, NOS2/MAC-3 double positive cells (n=10-16 per group). **(*C*)** Quantification of relative foam cell numbers (top) and size (bottom) in specified chimeric mice; *Trib1*^***mWT***^**→***ApoE*^*-/-*^ and *Trib1*^***mTg***^**→***ApoE*^*-/-*^ mice. N = 10-16, per group. (**D**) Representative image of aortic sinus lesion (scale: 30µm) from a *Trib1*^*m*WT^ *and Trib1*^*m*Tg^ mice injected with PCSK9 (n=9-11, per group, with arrows highlighting foam cells. Quantification of relative MAC-3 staining, foam cell numbers and size (n=6-7 per group). **(*E*)** Correlation between (1^st^ panel) foam cell number and MAC-3+ staining in plaque of aortic sinus lesions of specified *Trib1*^***mTg***^**→***ApoE*^*-/-*^ chimeric mice; (2^nd^ panel) foam cell size (y axis) and MAC-3 staining and (3^rd^ panel) plasma cholesterol levels (y axis) and macrophage staining. MAC-3+ immune-reactive area expressed as percentage (%) of total lesion area in aortic sinus. R^2^= Pearson correlation coefficient. In ***B* - *D***, data (mean ± SEM) are expressed relative to wild-type (WT). Significance was determined by one-way ANOVA (***B***) or student’s t-test (***C, D***). \**P* <0.05, \*\**P* <0.01, \*\*\**P* <0.001

In the *Trib1*^mTg^**→***ApoE*^-/-^ chimeras, there was a stronger correlation between the mean foam cell size and the percentage of aortic sinus stained by MAC3 than between MAC3^+^ staining and foam cell numbers (**Fig. 3E**) and, we could not ascribe the observed increase in plaque-foam cell numbers on the effects of m*Trib1* expression on blood cholesterol levels (**Fig 3E**), HDL-C levels or the non-significant rise in LDL-C (**Fig. S3**). In fact, while the *Trib1*^mWT^**→***ApoE*^-/-^ chimeras with the highest plasma cholesterol, HDL-C and LDL-C concentrations had the lowest amount of MAC3^+^ staining in their aortic sinus lesions, the inverse was true for the *Trib1*^mTg^**→***ApoE*^-/-^ chimeras (**Fig. 3E**, third panel**; Fig. S3**). Thus, collectively, these data indicate that increased macrophage lipid uptake/storage was the prominent driving force for the observed foam cell expansion besetting the early-stage of the atherosclerotic process in these and the *Trib1*^mTg^*-Pcsk9* mice.

### Myeloid-TRIB1 Expression Induces OLR1 Expression in Both Mouse and Man

To identify potential cellular mechanisms by which m*Trib1* enhances foam cell expansion, we analysed the gene expression characteristics of human *TRIB1*^High^ monocytes and *TRIB1*^High^ monocyte derived macrophages (MDM) using the microarray RNA data produced in the Cardiogenics Transcriptomic Study (25). In this dataset, involving samples from 758 individuals, *TRIB1* RNA levels were on average higher in monocytes than in MDMs (**Fig. 4A**, top panel) but, as is evident from the analyses of RNA levels in 596 paired samples, there was no correlation between *TRIB1* RNA levels in these two cell types (**Fig. 4A**, bottom panel). Moreover, genes differentially expressed in *TRIB1*^High^ versus *TRIB1*^Low^ monocytes were enriched for different sets of ‘DAVID’ Gene Ontology cluster terms (**Tables S1, S2**) than those characterising the more lipid-based transcriptome of *TRIB1*^High^ MDM (**Fig 4B, C, Tables S3**).

**Fig. 4.**
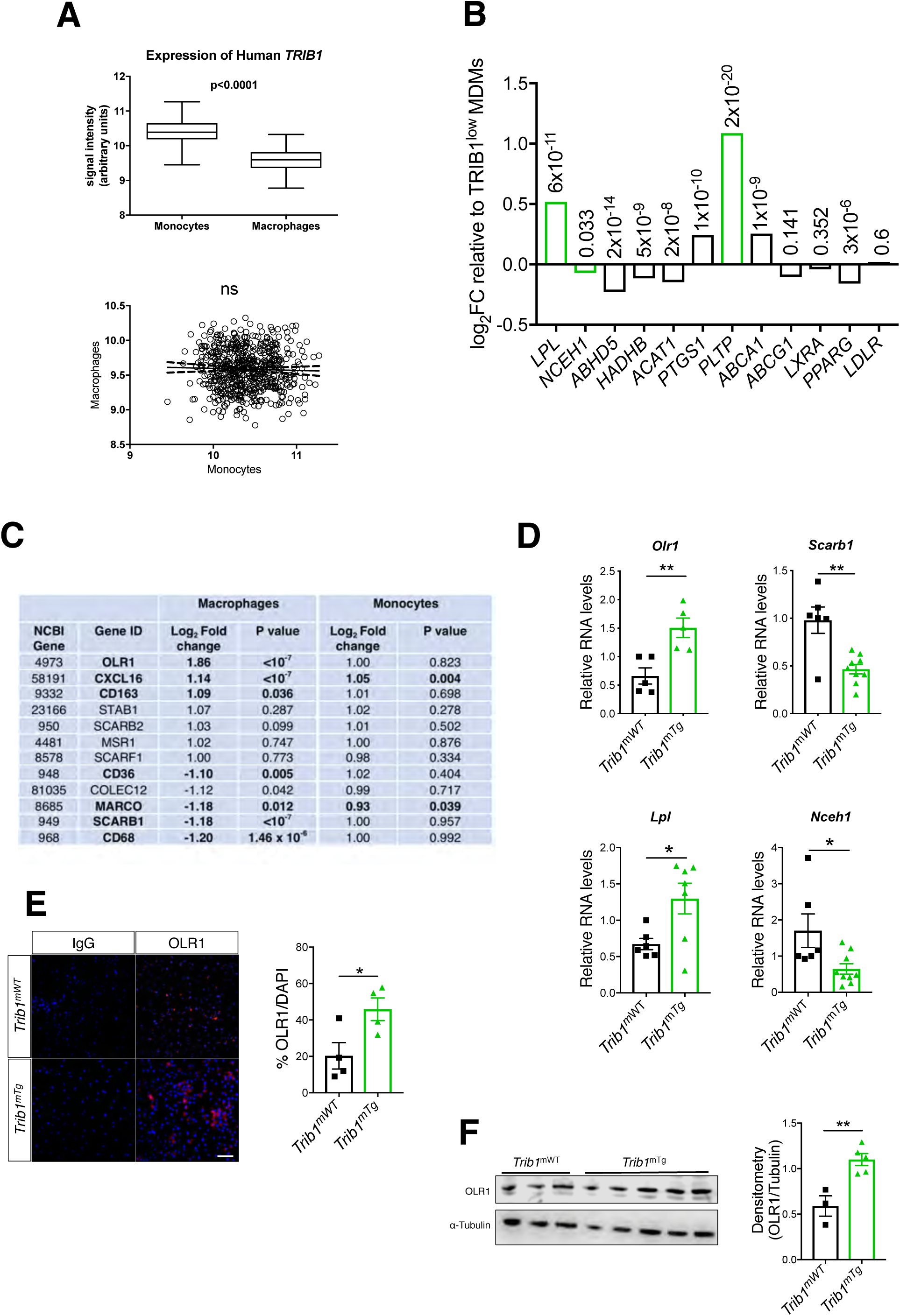
Myeloid TRIB1 Expression Induces Reciprocal Changes in oxLDL- and HDL-Receptor Expression in Human and Mouse Macrophages. *(A)* Comparison of *TRIB1* RNA levels in monocytes and monocyte-derived macrophages (MDMs) in participants of the Cardiogenics Transcriptomic Study (25). The correlation (R^2^<0.001, p=0.47) was performed on 596 paired monocyte and MDM samples. **(*B*)** MDM (n=596) and monocytes (n = 758) were ranked according to *TRIB1* RNA levels. Data represent log2-fold changes (FC) in RNA levels of specified genes in *TRIB1*^High^ (n=149) versus *TRIB1*^Low^ (n=149) MDM, with associated P values provided above the bars. Green bars indicate concordant changes in transcript levels of representative genes in human *TRIB1*^High^ MDMs and *Trib1*^mTg^ BMDMs. **(*C*)** FC and associated P values for differential expression of specified RNAs encoding representative scavenger receptors, including CD36, which mediates (ox)-phospholipid and long chain fatty acid uptake (37), the acetylated-LDL scavenging receptor(38) and Macrophage Scavenger Receptor (38). Comparisons are between *TRIB1*^HIGH^ (n=149) versus *TRIB1*^LOW^ (n=149) MDMs and between *TRIB1*^HIGH^ (n=191) versus *TRIB1*^LOW^ (n=191) monocytes. **(D)** RT-qPCR quantification of RNA levels in bone marrow derived macrophages (BMDM) prepared from specified mice (n= 5-9 per group mean ± SEM). n=5 **(E)** Immunocytochemistry of non-polarised *Trib1*^***mWT***^ and *Trib1*^***mTg***^ BMDMs. OLR1 (red), nuclei counterstained with DAPI (blue). Scale: 50µm. Quantification performed on BMDMs prepared from 4-5 mice per group. **(*F*)** Western blot analysis of OLR1 in *Trib1*^mWT^ and *Trib1*^mTg^ BMDMs (n=3-5 per group). In ***D*** - **F**, significance was determined by student’s t-test, \**P* <0.05, \*\**P*<0.01

The Cardiogenics Transcriptomic data strongly suggested that the m*TRIB1*-induced foam cell phenotype stemmed from increased oxLDL uptake rather changes in LDLR and scavenger receptor class B type 1 (which mediates selective HDL-cholesterol uptake and efferocytosis (18)) expression (**Fig 4C**) or reductions in ABCG1- and ABCA1-mediated cholesterol efflux (**Fig 4B**). Notably, *OLR1* was the fourth most differentially expressed gene in the *TRIB1*^High^ human MDMs and the most highly altered scavenger receptor in these cells (**Fig 4C**). We therefore examined the effect of m*Trib1* transgene expression on this oxLDL receptor. This revealed that *Trib1*^mTg^ BMDMs contained more *Olr1* RNA but fewer *Scarb1* transcripts (**Fig 4D**) than their *Trib1*^mWT^ counterparts, indicating that the increased numbers of *OLR1* and reduced numbers of *SCARB1* transcripts in human *TRIB1*^High^ MDMs are causally related to the increased number of *TRIB1* transcripts in these cells. The reciprocal relationship between *OLR1* and *SCARB1* RNA levels in *TRIB1*^High^ MDMs was also recapitulated in IFNγ/LPS, IL4-(**Fig. S5A**) and fatty acid-polarised MDM samples but not in HDL-polarized MDMs (**Fig. S5B, Table S6**). Rather HDL-polarised MDMs contained *OLR1* and *SCARB1* RNA levels indistinguishable from those of non-polarized MDMs (**Fig S5A, B, Table S6)** Finally, to substantiate the evidence for causal TRIB1 involvement in OLR1 expression we stained BMDMs for OLR1 protein. OLR1 was detected in twice as many *Trib1*^mTg^ cells than their wild-type counterparts (**Fig. 4E**), consistent with the Western Blotting analysis of whole BMDM cell lysates (**Fig. 4F**).

### m*Trib1*-Induced OLR1 Expression in Plaque Macrophages Increases Atherosclerotic Burden

To corroborate the evidence for causal mOLR1 involvement in plaque-resident macrophage foam cells expansion, we quantified OLR1 expression in the aortic sinus lesions of our mouse models using an OLR1-antibody that recognizes the cell-surface expressed form of this oxLDL receptor, as well as the proteolytically cleaved (soluble) extracellular form (9). The antibody detected OLR1 in MAC3+ cells and in acellular areas of the mouse aortic sinus lesions, including in regions adjacent to plaque-macrophages (**Fig 5A**). Moreover, as expected from the known expression and regulation of this scavenger receptor in endothelial cells (9), significant amounts of OLR1 was also detected in the non-macrophage (i.e. MAC3-) cell population at the plaque surface of the more hyperlipidemic model of human atherosclerosis (**Fig S3, Fig 5A**). Finally, confirming the causal involvement of mTRIB1 in mOLR1 expression (**Fig. 4D-F**), the anti-OLR1 antibody detected OLR1 in more of the plaque macrophages of *Trib1*^mTg^**→***ApoE*^-/-^ mice than in those of the *Trib1*^mWT^**→***ApoE*^-/-^ control animals (33.89 ± 6.56% vs. 16.18 ± 4.05%); a result which was replicated in the *Trib1*^mTg^-*Pcsk9* and *Trib1*^mWT^-*Pcsk9* mice (29.35 ± 7.42% vs. 10.38 ± 3.36%) (**Fig 5A**).

**Figure 5:**
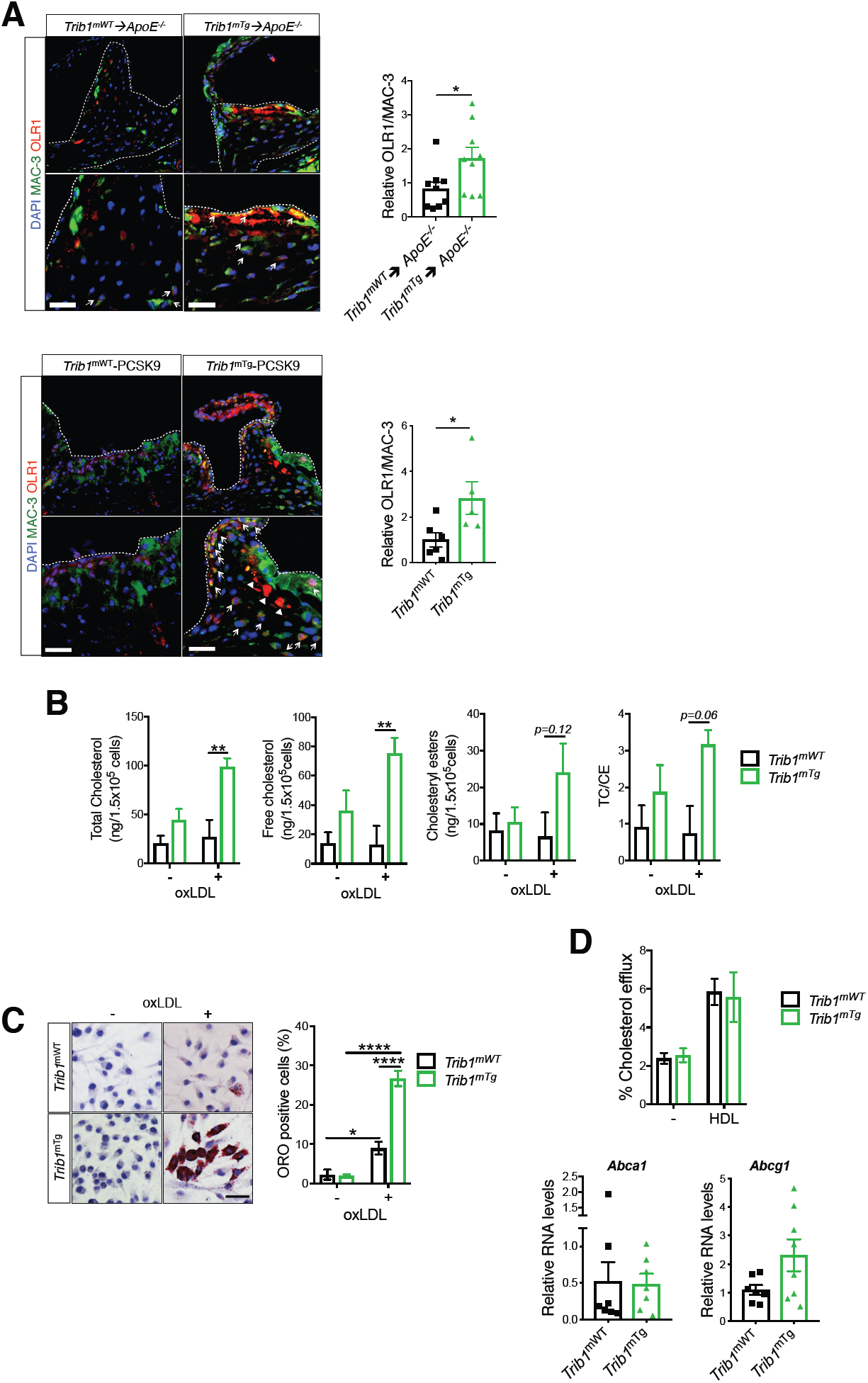
Myeloid-Trib1 Increases Cholesterol Uptake and Neutral Lipid Accumulation. **(*A*)** Representative images (Scale: 50µm) of aortic sinus lesions from specified mice and enlarged images. Dashed lines indicate boundaries of lesions. MAC-3 (green), OLR1 (red), nuclei counterstained with DAPI (blue). Arrows indicate OLR1-positive macrophages. Arrowheads indicate assumed a-cellular OLR1. Quantification: *Trib1*^***mWT***^**→***ApoE*^*-/-*^ and *Trib1*^***mTg***^**→***ApoE*^*-/-*^ chimeras, 9 per group; m*Trib1*-PCSK9, 5-6 per group (mean ± SEM). **(*B*)** Intracellular total cholesterol (TC) and cholesteryl esters (CE) contents of *Trib1*^mTg^ and *Trib1*^mWT^ bone marrow cells differentiated into macrophages and incubated with 25µg/ml of oxLDL for 24 hours (n=4 per group). **(*C*)** Representative image of BMDMs stained with Oil Red O (Scale: 50µm). Quantification was performed on three fields of view per sample. **(*D*)** Quantification of cholesterol efflux from cholesterol-loaded BMDMs to human HDL (n=8-9 per group). RT-qPCR quantification of *Abca1* and *Abcg1* RNA in non-polarised BMDMs prepared from specified mice (n= 7-8 per group). Data are mean ± SEM. Significance determined by student’s t-test (***A, D;*** *bottom panel*) or two-way ANOVA with Sidak’s multiple comparisons post-test (***B-D***) \**P* <0.05, \*\**P* <0.01, \*\*\*\**P*<0.001.

To mechanistically validate the contribution of *mTrib1*-induced Olr1 expression to foam cell expansion, we incubated non-polarised BMDMs with oxLDL for 24 h. This led to marked rises in the intracellular levels of both total cholesterol (2.71 ± 0.24-fold, P=0.0091) and unesterified cholesterol (5.81 ± 0.83-fold, P=0.0049) in the *Trib1*^mTg^ BMDMs but not in their WT-counterparts (**Fig. 5*B***). Additionally, as judged by Oil Red O staining, oxLDL transformed nearly three times as many *Trib1*^mTg^ BMDMs into foam cells than *Trib1*^mWT^ cells (**Fig. 5*C***, P <0.0001), as evidenced by the very visible increase in neutral lipid accumulation in these cells upon exposure to oxLDL. In contrast to the profound effects of oxLDL on cholesterol accumulation in un-polarised *Trib1*^mTg^ BMDMs, we observed no impairment of HDL-mediated cholesterol efflux (**Fig. 5*D***), consistent with the observation that neither *Abca1* nor *Abcg1* RNA levels are reduced in these BMDMs (**Fig 5D**) and, that the *Trib1*^mTg^**→***ApoE*^-/-^ chimeras have higher, rather than lower, HDL-C levels than their wild-type peers (**Fig S3**). Thus, collectively, our results suggest that *Trib1*-induced foam cell expansion in early-stage atherosclerotic plaque arises from increased cholesterol/neutral lipid uptake and retention rather than reduced HDL-mediated cholesterol efflux.

## Discussion

Despite the success in establishing that hepatic *Trib1* expression affects the regulation of multiple cellular processes modulating blood cholesterol and triglyceride levels (16), the influence of global-knockout of *Trib1* on shaping the phenotype of macrophages (17), and the finding that variants at the *TRIB1* locus are associated with and increased CHD risk (14, 15), the contribution of Tribbles-1 on atherogenesis remains to be addressed. Herein, we demonstrate that there is a wide distribution of *TRIB1* RNA levels in human MDMs and, that genetically engineered changes in m*Trib1* expression in mouse models of early-stage human atherosclerosis markedly affect the size of developing plaques and the morphological and functional properties of plaque-macrophages (**Fig. 6**). In summary, we have confirmed the pro-atherogenic impact of myeloid TRIB1 in two distinct *in vivo* models of human atherosclerosis.

**Fig. 6.**
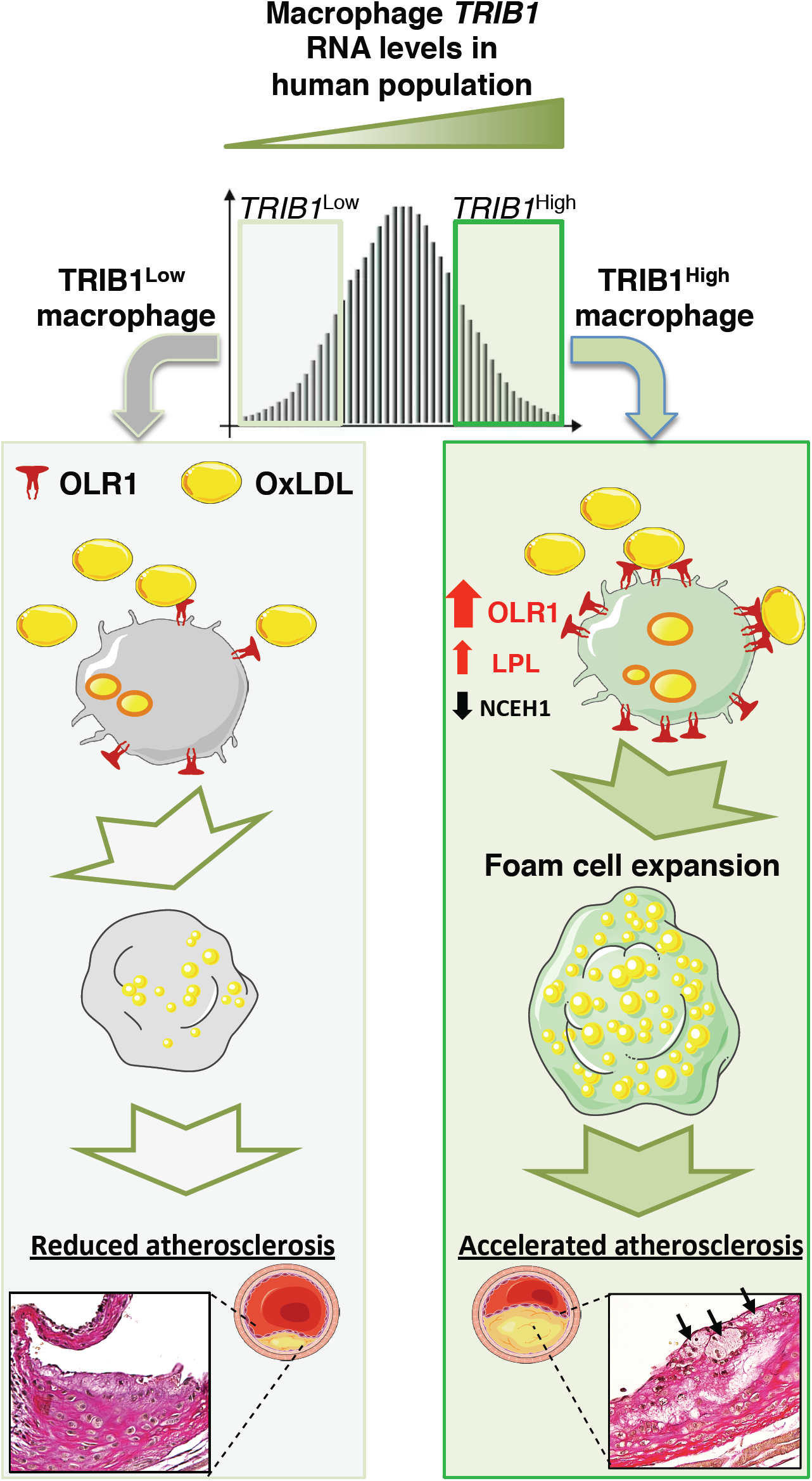
Model Summarising the Proposed Effects of Differences in mTRIB1 Expression on Foam Cell Expansion in Early-stage Atherosclerosis. Factors up-regulating m*TRIB1* expression in human monocyte-derived macrophages and plaque-macrophages increase cholesterol and neutral lipid uptake, with no compensatory rise in cholesterol-efflux. Schematic recognises that increased OLR1 expression increases the probability of this scavenger receptor assembling as a hexamer made up of three homodimers on the macrophage cell surface and that this configuration leads to a marked increase in its affinity for oxLDL (9) and, hence OLR1-mediated uptake of oxLDL lipids. In the setting of no compensatory rise in HDL-mediated cholesterol efflux, accelerated foam cell expansion and increased atheroma burden ensue, highlighting the therapeutic potential of inhibiting macrophage Tribbles 1 expression to block the gene expression changes that promote macrophage cholesterol and cholesteryl ester accumulation and prevent increased hydrolysis of accumulated cholesteryl-ester and thus, the up-regulation of the reverse cholesterol transport pathway to mediate the removal of cholesterol from the arterial wall.

Recent studies have established that oxLDL accumulates steadily in both early-(growing) and mature-(yellow plaque without a necrotic core) stage human coronary plaques but that in more advanced vulnerable plaques (yellow plaques with a necrotic core) this lipoprotein is removed either by metabolism or replacement with other substances, including cell debris (2). This *in vivo* data dove-tails well with the early *in vitro* work which showed that while oxLDL promotes macrophage growth and survival in a dose-dependent manner, beyond a certain lipid concentration cell death ensues, albeit by an unknown mechanism (10). Thus, a critical question to consider is whether m*TRIB1*-induced OLR1 expression serves an (athero-) protective role in the early stage of human atherosclerosis, for example, by reducing the exposure of plaque-resident vascular cells (where this disease is initiated) to oxLDL. This lipoprotein is a well-described activator of endothelial cell OLR1 expression with the totality of the data indicating that this activation culminates in arterial endothelium dysfunction (9). In the experiments reported here we provide evidence that, m*Trib1* transgene expression reduces vascular cell exposure to oxLDL given that it increased the size and lipid contents of plaque-resident foam cells in two independent models of early-stage atherosclerosis by increasing mOLR1 expression and oxLDL uptake. Notably, in the less hyperlipidaemic of these two transgenic models, we also could discern a very strong positive correlation between plaque-macrophage numbers and plasma cholesterol/LDL-C levels, implying that in early-stage atherosclerosis m*Trib1*^High^ macrophages (in marked contrast to Trib1-deficient macrophages) are uniquely equipped to increase plaque-macrophage numbers in response to lipid excess.

Based on the strong evidence of a causal link between hyperlipidaemia and CHD (26) and, the demonstration that ablating hepatic *TRIB1* expression increased plasma levels of cholesterol, LDL-C and of triglyceride (16), the implication was, which seemed entirely consistent with GWAS results (14, 15) that increasing *TRIB1* would be athero-protective. Silencing *TRIB1* expression in macrophages, however, turns out to be athero-protective, as judged by analyses of the aortas and aortic sinuses from *Trib1*^mKO^**→***ApoE*^-/-^ chimeric mice, after a 12 weeks Western dietary regime. This counter-intuitive result, which fits well with our *in vitro* and *in vivo* analysis of the consequences of m*Trib1*^High^ expression on foam cell expansion, suggests moving forward that developing a therapy to specifically silence *Trib1* expression in macrophages would provide clinical benefit beyond that of lipid-lowering medications, although full realization of this benefit may require its adoption at an early-stage of atherogenesis. More generally, our study also demonstrates that mechanistic probing of GWAS signals is not only warranted, but critical, to identify both the totality and directionality of disease risk factors, even when, as was the case for *TRIB1*, the association between a genetic variant(s), disease risk and a major disease-risk factor (plasma lipids) appeared congruent. Hence, we acknowledge that a limitation of the current study is that we have not addressed whether changes in vascular cell *TRIB1* expression might affect early-stage atherosclerosis development and, whether the GWAS-CHD signal at the *TRIB1* locus reflects that in these cells (and hepatocytes) TRIB1 serves an athero-protective role, in contrast to the situation in plaque-resident macrophages where it induces foam cell expansion.

One of the most striking outcomes arising from roughly doubling *Trib1* expression in macrophages was the increase in foam cell size that developed in both of our models of human atherosclerosis. Our results indicate that this was driven by the failure of these cells to increase HDL-mediated cholesterol efflux in response to an up-regulation of cholesterol and fatty acid uptake. Additionally, this up-regulation was attributable to increased OLR1 RNA and protein expression, consistent with the *in vitro* studies of Lazar and coworkers (27) which demonstrated that Rosiglitazone-induction of *Olr1* expression in adipocytes increased oxLDL and palmitate uptake and cellular cholesterol levels. Likewise, when Steinbrecher and colleagues (28) increased macrophage *OLR1* expression via lysophosphatidylcholine stimulation, oxLDL uptake was also induced, prompting them to introduce the concept that macrophages within atheromas may be quiescent with respect to oxLDL uptake until *Olr1* expression is induced (28). Here, we validate this concept by demonstrating overexpressing *mTrib1*-led directly to an increase in OLR1 protein in plaque-resident macrophages, alongside their increased lipid contents and size. Our data also show that by doubling m*Trib1* expression, we removed the requirement for an external stimulus to up-regulate *OLR1* expression and the consequent clinical sequelae. In fact, the phenotype observed in response to this rise in *mTrib1* expression fits well with earlier computational modelling studies that indicated that relatively modest changes in TRIB1 would have a major impact on the activity of MAPK pathways, thus defining the concentration of these proteins is critical for shaping cell function (29).

Thus, from the therapeutic standpoint our results reveal that targeting TRIB1 expression could, in addition to beneficially affecting OLR1 expression, moderate CHD pathogenesis by simultaneously altering in favourable directions the expression of a number of other disease-promoting genes affected by changes in TRIB1 expression. These would include, for example, beneficially increasing NCEH1 expression to reduce the release of pro-inflammatory cytokines from plaque-resident macrophages (30), while reducing the expression of LPL to help reduce the retention of LDL in the artery wall during early-stage atherosclerosis (7), as well as excessive accumulation of cholesteryl ester and triglycerides within macrophage foam cells (8). Whether mTRIB1 modulates atheroma regression (31, 32) and late-stage atherosclerotic plaque stability by orchestrating a coordinated response to the lipid and inflammatory challenges encountered by plaque-resident macrophages/foam cells in advanced stage atherosclerosis now requires investigation.

## Materials and Methods

### Human samples

Human tissue and blood samples were collected under protocols approved by the University of Sheffield Research Ethics Committee and Sheffield Teaching Hospitals Trust Review Board (Ref. STH 16346, SMBRER310) and in accordance with the Declaration of Helsinki. All participants gave written informed consent. Human coronary arteries obtained from explanted hearts were fixed in 10% (v/v) formalin and embedded in paraffin wax. Antigen retrieval was performed with trypsin for 15 mins at RT (#MP-955-K6, A Menarini Diagnostics, UK). Sections were incubated with mouse anti-human CD68 (#M0814, Clone KP1, Dako) antibody (1:100 dilution) and rabbit-anti-human TRIB1 (#09-126, Millipore, UK) antibody (1:100) and the appropriate secondary antibodies, biotinylated horse anti-mouse secondary antibody and goat anti-rabbit secondary (#BA-2000, #BA-1000, Vector Laboratories) antibody (both 1:200 dilution); and the detection reagents, Elite Mouse ABC HRP (#PK-6100, Vector Laboratories), 3, 3’-Diaminobenzidine (SIGMA*FAST*™, D4293, Sigma), rabbit ABC-Alkaline phosphatase and Vector Red Alkaline phosphatase substrate kit (AK-5000, #SK-5100, Vector laboratories). Sections were counterstained with haematoxylin and mounted with DPX mountant (#44581, Sigma-Aldrich).

### Mice and creation of murine models of myeloid-specific *Trib1* expression

Mice were handled in accordance with UK legislation (1986) Animals (Scientific Procedures) Act. Mouse experiments were approved by the University of Sheffield Project Review Committee and carried out under a UK Home Office Project Licence (70/7992). All mice used were congenic on a C57BL/6J background (N17) and were housed in a controlled environment with a 12-hour light/dark cycle, at 22°C in Optimice individually ventilated cages (Animal Care Systems) and given free access to a standard chow diet (#2918; Harlan Teklad) and water. A KOMP repository embryonic stem (ES) cell clone containing loxP sites flanking exon 2 of *Trib1* (EPD0099_5_D04) was used to generate a floxed *Trib1* allele. The clone was genotyped to validate its authenticity, injected into C57BL/6J blastocysts and transferred to pseudo-pregnant recipient females (Geneta). Resulting chimeras were mated with a FLP-deleter strain maintained on a C57BL/6J background (Geneta) and floxed-*Trib1* mice generated. *Trib1*^mKO^ mice were generated by crossing floxed-mice with Lys2-cre-recombinase transgenic mice (https://www.jax.org/strain/004781), excising all but the first 120 amino acids of TRIB1. Murine *Trib1* cDNA was introduced into the previously described, pROSA26, loxP-flanked STOP and Frt-flanked IRES-eGFP targeting construct (33) and then inserted into the ubiquitously expressed *Rosa26* locus of Bruce4 (C57BL/6 origin) mouse ES cells by homologous recombination. Correct integration was confirmed by Southern blotting. *Trib*^1mTg^ mice were generated by crossing floxed mice with the Lys2-cre recombinase transgenic mice described above. Mice were genotyped by PCR amplification of ear-clip samples. *Trib1 fl/fl* x *Lyz2Cre* and *Rosa26*.*Trib1* x *Lyz2Cre* were genotyped for the presence of *Lyz2Cre* and for either *Trib1 fl/fl* or *Rosa26*.*Trib1* using three primer sets (**Table S4**). For the atherosclerosis experiments mice were fed a Western (21% fat, 0.2% cholesterol) diet (829100; Special Diet Services, Braintree, UK) for 12 weeks.

#### mTrib1 RNA quantification

Bone marrow derived macrophages (BMDM) were cultured for five days in complete medium: DMEM (BE12-604F, Lonza) medium containing 10% (v/v) L929-conditioned medium, 10% (v/v) ultra-low endotoxin FBS (S1860-500, BioWest, USA) and 100 U/ml Penicillin and 100ng/ml Streptomycin (15140-122, Gibco). Total RNA was isolated using ReliaPrep™ kit (#Z6011, Promega) and reverse transcribed into cDNA using iScript cDNA synthesis kit (#1708890, Bio-Rad). *Trib1* was quantified using the TaqMan® assay Mm00454875, which amplifies a 99-nucleotide amplicon comprising exon 2 and 3 sequences. *mGFP quantification*. GFP positive cells in freshly purified peripheral blood monocytes were isolated by positive selection using magnetic MicroBeads conjugated with F4/80 (#130-110-443, Miltenyi Biotec) and CD115 (#130-096-354, Miltenyi Biotec) using a modified version of the protocol described by Houthuys *et al* (34). Inflammation was induced by injecting PBS containing thioglycollate into the peritoneal cavity and isolating the infiltrating cells, as described (35). The percentages of GFP-positive cells in blood and bone marrow cells (from femurs and tibiae) were determined using mice sacrificed humanely by cervical dislocation. Red blood cells were lysed using RBC lysis buffer (eBioscience, 00-4300-54). Approximately 10^6^ cells/sample were re-suspended in PBS and dead cells removed using amine-reactive dye NIR Zombie (1:500 dilution in the antibody master mix; #423105, BioLegend). Live cells were stained with the following cell surface marker specific antibodies, at 0.1μg/ml each in 100μl total volume: AF647-conjugated anti-human/mouse CD11b (#101220, BioLegend), PE-conjugated anti-mouse Ly6C (#101220, BioLegend) and PE/Cy7-conjugated anti-mouse Ly6G (#127617, BioLegend). Cells were sorted on an LSR II Cytometer (BD Bioscience) equipped with 355nm, 405nm, 488nm and 633nm excitation lasers. Quantifications were performed with FlowJo software.

#### Blood Counts

Blood counts of heparinized blood, obtained via cardiac puncture, were determined using a Sysmex KX-21N quantitative automated haematology analyser.

#### Histological analysis

Adipose tissue and liver samples were paraffin-embedded. Cross-sections were stained with Haemotoxylin and Eosin or incubated with F4/80 (1:50; #565409 (Clone T45-2342), BD Pharmingen), followed by biotinylated rabbit anti-rat antibody (1:200, Vector Laboratories, UK) and PE-Streptavidin (1:20, #405203, Biolegend). They were counterstained with DAPI (#P36931, Invitrogen) and mounted with ProLong® Gold anti-fade mountant. Mean adipocyte sizes were determined using NIS-Elements software (Nikon Instruments, UK) by measuring at least 15 cells per field of view and, three fields of view per mouse. Frozen spleen sections were stained with F4/80 (PE-rat anti-mouse #123019 (Clone BM8, BioLegend) or CD206-(Alexa Fluor-647 rat anti-mouse #321116 (Clone 15-2, BioLegend) conjugated antibodies (1:200), counterstained with DAPI and mounted with Aquamount (Thermo Fisher Scientific). Fluorescent images were captured using an inverted wide-field fluorescence microscope (Leica AF6000).

### Models of early-stage human atherosclerosis

For the bone marrow transplantation model, 12-13 week old mixed gender donor mice were sacrificed humanely by cervical dislocation. Femur/tibae bone marrow cells were isolated and purified by standard methods and re-suspended in Hank’s buffered salt solution (HBSS, without phenol red, #14175053, ThermoFisher) containing 10% (v/v) foetal calf serum. Donor cells (2-4 × 10^6^) were transplanted via tail-vein injection to randomly allocated 12-13 week old male *ApoE*^-/-^ recipient mice who were lethally irradiated with 11 Grays in two doses (5.5 Gy on two occasions separated by 4 hours) on the day of the transplantation. Bone marrow transplant experiments were undertaken in two waves, one for each m*Trib1* model and respective WT control. During the seven weeks post-transplant recovery period, the chimeras were fed a standard chow (#2918; Harlan Teklad) diet and then switched to Western diet (21% fat, 0.2% cholesterol, 829100; Special Diet Services, Braintree, UK) for 12 weeks. Chimera mice were given sterile acidified drinking water (1.1% (v/v) HCl) until the end of the procedure.

#### PCSK9 model

An adeno-associated virus-based vector (rAAV8) that supports transport of the m*PCSK9*^*D377Y*^ gene to the liver was purchased from UNC GTC Vector Core (Chapel Hill, NC). The mice received 6.1 × 10^11^ viral particles via a single tail vein injection. Following 7 days recovery, they were transferred to the Western diet (829100, Special Diet from Braintree, UK) for 12 weeks.

#### Lipid measurements

Fasting, plasma total cholesterol, HDL-C, triglycerides and glucose levels were measured on a Roche Cobas 8000 modular analyser. LDL-C was estimated by the Friedewald equation. Additionally, colorimetric assays (Cholesterol Quantification kit, #MAK043, Sigma Aldrich; Triglyceride Assay kit, #ab65336, Abcam) were used.

### Atherosclerotic Plaque Assessment

Mice were perfused through the heart with PBS and then with 10% (w/v) neutral buffered formalin. The aorta was segmented at the diaphragm and at the top of the aortic arch. Following careful removal of surrounding extraneous fat and connective tissue, it was dissected longitudinally and fixed in 4% (w/v) paraformaldehyde. Dissected aortas were stained with Oil Red O (60% (v/v) in isopropanol) and pinned onto wax. Images were taken with a macroscopic CCD camera and analysed by computer-assisted image analysis (NIS-Elements software, Nikon Instruments, UK). Aortic sinus samples were obtained by excising the heart and transecting parallel to the atria. Following fixation in 10% formalin (v/v) buffered saline for at least 24 hours, samples were serially cut (at 7µm intervals) from the valve leaflets until the beginning of the aorta.

#### Immunohistochemistry

Aortic sinus sections were dewaxed and rehydrated. Heat-mediated antigen retrieval was performed with 10mM sodium citrate and non-specific staining reduced by incubation in 5% (v/v) goat serum (#G9023, Sigma Aldrich) for 30 mins at RT. Primary antibodies diluted as appropriate were: the rat anti-mouse MAC-3 antibody Clone M3/84 (1:100 dilution, BD Pharmingen), rabbit polyclonal NOS2 antibody (#ab15323; Abcam, UK 1:100), rabbit anti-mouse YM1 polyclonal antibody (#ab93034; Abcam, UK 1:100) and or rabbit polyclonal OLR1 (#ab203246; Abcam, UK, 1:100). The secondary antibodies were biotinylated rabbit anti-rat secondary antibody (1:200, Vector Laboratories BA-4000), goat anti-rat DyLight®488 goat anti-rat DyLight®488 (#GtxRt-003488NHSX, ImmunoReagents Inc.) and goat anti-rabbit DyLight®550 (#GtxRb-003-D550NHSX, ImmunoReagents Inc.). Rabbit anti-rat conjugated biotinylated antibody was visualised with the Vectastain ABC-HRP complex (PK-6100, Vector Laboratories). Sections were counterstained with Carazzi’s haematoxylin. The number and area of foam cells (MAC-3^+^ macrophages containing a vacuolated cytoplasm) were quantified using NIS-Elements software (Nikon, UK). Fluorescent images were captured using an inverted wide-field fluorescence microscope (Leica AF6000). Quantification of YM1, NOS2 and OLR1 in the aortic sinus lesions were confined to macrophage (MAC3+) cells only and analysed using ImageJ.

#### Clinical Grading

The atherosclerotic burden of the atheroma in aortic sinus samples was graded according to the Stary system (e.g. 1 = presence of macrophage foam cells, 2 = presence of intracellular lipid accumulation, 3= presence of extracellular lipid pools) (36) using a light microscope (Zeiss Axiophot, Carl Zeiss, Jena, Germany) and Imageaccess^©^. Slides were randomized and the cardiac pathologist was fully blinded to sample origin.

### Western blotting

Twenty microgram (µg) of total protein lysates were size-fractionated on 4-12% NuPAGE Bis-Tris gel (#NP0321, Invitrogen). OLR1 was detected by incubation with a rabbit anti-mouse OLR1 antibody (1:500, #ab203246; Abcam), followed by HRP conjugated goat-anti-rabbit IgG (#P0448, Dako (1:1000). LDLR was detected by incubation with a rabbit polyclonal antibody (1:1000, (#3839-100, BioVision, USA) 4°C overnight followed by HRP conjugated goat-anti-rabbit IgG (#P0448, Dako (1:1000). α-Tubulin was used as a housekeeping control (#sc-32293, Santa Cruz). Detection of immuno-reactive products was performed using 1:1 ECL reagent (#RPN2235, GE Healthcare) and a C-DiGit® Blot scanner (Model 3600, LI-COR). Densitometry was performed with Image Studio™ Lite software (LI-COR).

### Gene expression studies

RNA from BMDM was prepared as described above. RT-qPCR was performed using primer sequences provided in **Table S5** and either SYBR-Green (PrecisionPLUS, Primer Design, UK) or a Taqman (Invitrogen) assay (*TRIB1*, Hs00921832; *OLR1* Hs01552593; *SCARB1*, Hs00969821, ThermoFisher). Values were normalised to either *Actb* (Mm02619580, Invitrogen) for mouse samples or *GAPDH* for human samples (Hs02786624), Invitrogen). Fold changes were calculated using the ΔΔCt method.

#### Microarray analyses

Details of the Cardiogenics Transcriptomic Study (CTS), which comprises transcriptomic data from isolated monocytes from 758 donors and matched monocyte derived macrophage (MDM) samples from 596 donors have been published (25). In brief, monocytes were isolated from whole blood using CD14 immuno-beads and cultured for 7 days with macrophage colony-stimulating factor (MCS-F) to generate macrophages. The top and bottom quartiles of *TRIB1-*expressing samples were defined respectively as *TRIB1*^High^ and *TRIB1*^Low^. Comparable sizes of differentially expressed gene lists (n=1842 monocytes; n= 2171, MDM) were obtained by using FDR adjusted p-values (i.e. q-values) of < 0.01 plus cut-off log-2 fold changes of > 0.071 (upregulated) and > −0.071 (down-regulated) for the MDM dataset. The <u>D</u>atabase for <u>A</u>nnotation, <u>V</u>isualization and <u>I</u>ntegrated <u>D</u>iscovery version 6.8 was used to identify for enrichment of functionally related Gene Ontology terms.

#### Gene expression in polarized human macrophages

Healthy human monocytes were isolated from whole blood by Ficoll-Paque PLUS (17144003, GE Healthcare) density centrifugation followed by magnetic selection with CD14 Human MicroBeads (130-050-201, Miltenyi Biotec). Monocytes were incubated for 7 days with RPMI-1640 (Gibco) supplemented with 100 ng/ml MCS-F (#300-25-100, PeproTech, UK), 10% FBS (Biowest), 1 mM glutamine (Invitrogen) and 1% penicillin/streptomycin (Gibco), followed by polarization with either 100 ng/ml LPS (581-007-L002, Enzo Life Sciences, USA) and 20 ng/ml IFN-γ (300-02, PeproTech, UK); 20 ng/ml IL-4 (200-04, PeproTech, UK) or 20 ng/ml IL-10 (#200-10, PeproTech, UK) for 24 h. RNA and RT-qPCR were performed, as described above using the primer sets listed in **Table S4.**

### Ox-LDL uptake and HDL-mediated cholesterol efflux

Assays were performed on BMDMs cultured for 5 days as described above. OLR1 was detected as described above. For the uptake assays, cells were incubated at 37^0^C for a further 24 h with human oxLDL (#5685-3557, Bio-Rad) at a concentration of 0 µg/ml and 25µg/ml. Total cell lipids were extracted using 7:11:01 (v/v) chloroform:isopropanol:IGEPAL® CA-630 (#I8896, Sigma-Aldrich), dried at 50°C and re-suspended in 120µl cholesterol assay buffer (#MAK043, Sigma-Aldrich). Total cholesterol, free cholesterol and cholesteryl esters were measured using a Cholesterol quantification kit (#MAK043, Sigma-Aldrich), according to the manufacturer’s instructions. Foam cells were assessed by Oil Red O (60% (v/v) in isopropanol) staining as described above. For the efflux assays, BMDMs were incubated for 24 hours in DMEM medium (BE12-604F, Lonza) supplemented with 0.2% (w/v) fatty acid free-BSA (#A8806, Sigma-Aldrich) and 2.5µM TopFluor® (Bodipy) cholesterol (Avanti® Polar Lipids, Inc. USA). The medium was removed and the cells washed with PBS and equilibrated for 18 hours in DMEM supplemented with 0.2% (w/v) fatty acid free BSA Efflux was measured after a 4-hour incubation period with 0µg/ml and 50µg/ml of human HDL (#5685-2004, BioRad). Supernatants were collected and the cells lysed with 1% (w/v) cholic acid (#C1129, Sigma Aldrich) in ethanol. Cholesterol efflux was calculated as the percentage of fluorescence (excitation 490nm, emission 520nm) in the cell medium at the end of the incubation period divided by the total fluorescence in the medium and cells.

### Statistics

All data are reported as mean ± SEM unless stated otherwise in the figure legend. Graphs were produced and analysed by GraphPad Prism software. Each data point represents a single mouse or human donor. P values <0.05 were considered significant.

## General

We would like to thank Dr Markus Ariaans, Mr Carl Wright and the BSU staff at the University of Sheffield for their technical support during the *in vivo* experiments; Ms Fiona Wright for tissue processing and histology; Dr Steven Haynes for mouse genotyping and Ms Laura Martínez-Campesino for assistance and logistical support.

## Author contributions

Designing research studies: TND, HLW, AHG, DR, CCS, SEF, EKT

Conducting experiments and acquiring data: JMJ, AA, RB, KB, TND, MT, SEF

Analyzing data: JMJ, RB, SH, SKS, KB, ZH, TND, HLW, AHG, CCS, SEF, EKT

Writing the manuscript: JMJ, HLW, AHG, SEF, CCS, EKT

## Supplementary Materials

**SFig. 1. Expected and observed numbers of 8-week old offspring with specified *Trib1* genotypes**

(**A)** The *Trib1* ‘KO-first’ targeting construct contains elements to produce a full-body *Trib1* deletion-null allele (tma1). No offspring with a tma1/tma1 genotype were identified from this cross involving two heterozygote parents. **(B)** The expected numbers of mice with each of the three theoretical genotypes (i.e. homozygous WT (+/+), heterozygous (+/tm1c) and homozygous (tm1c/tm1c) from crossing two heterozygote parents were obtained. **(C)** Homozygous conditional-ready (i.e. floxed) *Trib1* null mice (i.e. tm1c/tm1c) were crossed with the heterozygous universal-Cre-recombinase mouse strain B6.Cg-Tg(UBC-cre/ERT2)1Ejb/J. N.B the fewer than expected mice homozygous for Tm1d-null *Trib1* allele. **(D)** Genotype distribution of offspring with the (tma1) *Trib1* deletion null allele on a mixed genetic (C57BL/6 × 129S9) background. **(E)** Representative gating strategy showing SSC-A and FSC-A plots of specified cell populations from blood and bone marrow **(F)** from 10-14 week old *Trib1*^mWT^ and *Trib1*^mTg^ mice (n=4). Live/dead cell discrimination was determined by Zombie NIR amine-reactive dye staining. CD11b^+^ cells were subdivided based on their expression of Ly6C (Y axis) and Ly6G (X-axis). Arrows indicate the fate of specific cell populations. **(G)** TRIB1-GFP expressing cells in CD11b^+^Ly6C^+^Ly6G^-^ (top left quadrants of (F)) and CD11b^+^Ly6C^-^Ly6G^+^ (bottom right quadrants of (f). (H, I) Quantification of GFP-positive cells within CD11b^+^Ly6C^+^Ly6G^-^ and CD11b^+^Ly6C^-^ Ly6G^+^ populations of blood and bone marrow cells (n=4, mean ± SEM).

**SFig. 2. *Trib1***^**mKO**^ **and *Trib1***^***mTg***^ **mice have normal tissue anatomy and F4/80+ macrophage numbers**

**(A)** Representative H&E staining of adipose tissue (visceral) cross-sections from 10-week old specified mice fed on chow diet. Middle panel, mean adipocyte area of samples (n=3-7, mean ± SEM); Left panel, F4/80+ macrophages contents of adipose tissue samples from specified mice (n=3-7, mean ± SEM). **(B)** Representative H&E staining of liver cross-sections and levels of F4/80+ macrophages in the liver (n=3-9, mean ± SEM); Scale, 20µm. **(C)** Representative IF staining of spleens from *Trib1*^mKO^, *Trib1*^mWT^and *Trib1*^mTg^ mice stained with F4/80 (red, left panels) and CD206 (green, right panels). Dotted lines indicate outlines of red and white splenic pulp. Scale: 50µm. ns: non-significant.

**SFig. 3. Plasma lipid levels of chimera and Pcsk9 mice.**

**(A)** Plasma lipid levels of specified chimera mice following seven weeks recovery and 12 weeks on Western Diet (n=6-12, mean ± SEM). **(B)** Total plasma cholesterol and triglyceride of mTrib1 Pcsk9 mice (n=7, mean ± SEM). **(C)** Correlations in specified mice between total plasma cholesterol (left panel), HDL-C (centre) and LDL-C (right panel) vs. oil Red O (top panels) and MAC-3^+^ immuno-reactive areas (bottom two panels), expressed as percentage (%) of total lesion area in aortic sinus. Data shows Pearson correlation co-efficient (R^2^) along with *P* value (n=9-10).

**SFig. 4. Atherosclerotic burden in m*Trib1*→ ApoE**^**-/-**^ **mice, clinical grading of lesions and presence of foam cells.**

**(A)** *en face* oil Red O staining of the thoracic aortas (n=6-15, mean ± SEM) and **(B)** lesion sizes in the aortic sinus (n=6-12, mean ± SEM) of *Trib1*^*mWT*^**→** ApoE^-/-^ versus *Trib1*^*mTg*^ **→** ApoE^-/-^ (left-hand panels) and *Trib1*^*mWT*^**→** ApoE^-/-^ versus *Trib1*^*mKO*^ **→** ApoE^-/-^ (right-hand panels) mice. Data are expressed as a percentage (%) of total surface area of the whole aorta. **(C)** Pathological grading of aortic sinus lesions assessing plaque fibrosis (left panel) and overall Stary Grade (e.g. 1 = presence of macrophage foam cells, 2 = presence of intracellular lipid accumulation, 3= presence of extracellular lipid pools) (centre panel) including featured of necrosis and haemorrhage. Multiple lesions per mouse (n=10-16, mean ± minimum and maximum) were scored indicate early stage lesions. Collagen content in the aortic sinus was quantified with Martius Scarlet Blue (right panel, n=10-16, mean ± SEM). (**D**) Representative image of aortic sinus lesion (scale: 30µm) from m*Trib1***→** ApoE^-/-^ mice, with arrows highlight foam cells. Significances were determined by student’s t-test (**A-B**) or one-way ANOVA **(C)**. Ns, non-significant, \**P*<0.05, \*\**P*<0.01.

**SFig. 5. Reciprocal regulation of OLR1 and SCARB1 RNA levels in polarised MDMs**

**(A)** *OLR1* and *SCARB1* RNA levels were quantified by RT-qPCR in human MDMs polarised with interferon γ + lipopolysaccharide (IFN γ + LPS), IL-4 and IL-10. Each data point represents data from one donor (n=8). Log2-fold changes (mean ± SEM) are plotted relative to non-polarised cells**. (B)** Fold-changes in levels of specified transcripts in human MDMs polarised with lauric acid (LA), linoleic acid (LIA), oleic and (OA) and HDL. Data were extracted from transcriptome analysis of Xue and colleagues (39).

## References and Notes

1. Rocha VZ, Libby P (2009) Obesity, inflammation, and atherosclerosis. Nat Rev Cardiol 6(6):399–409.

2. Uchida Y (2017) Recent Advances in Fluorescent Angioscopy for Molecular Imaging of Human Atherosclerotic Coronary Plaque. J Atheroscler Thromb 24(6):539–551.

3. Moore KJ, Sheedy FJ, Fisher EA (2013) Macrophages in atherosclerosis: a dynamic balance. Nat Rev Immunol 13(10):709–721.

4. Skålén K et al. (2002) Subendothelial retention of atherogenic lipoproteins in early atherosclerosis. Nature 417(6890):750–754.

5. Schrijvers DM, De Meyer GR, Kockx MM, Herman AG, Martinet W (2005) Phagocytosis of apoptotic cells by macrophages is impaired in atherosclerosis. Arterioscler Thromb Vasc Biol 25(6):1256–1261.

6. Tabas I (2010) Macrophage death and defective inflammation resolution in atherosclerosis. Nat Rev Immunol 10(1):36–46.

7. Gustafsson M et al. (2007) Retention of low-density lipoprotein in atherosclerotic lesions of the mouse: evidence for a role of lipoprotein lipase. Circ Res 101(8):777–783.

8. Takahashi M et al. (2013) Macrophage lipoprotein lipase modulates the development of atherosclerosis but not adiposity. J Lipid Res 54(4):1124–1134.

9. Hofmann A, Brunssen C, Morawietz H (2017) Contribution of lectin-like oxidized low-density lipoprotein receptor-1 and LOX-1 modulating compounds to vascular diseases. Vascul Pharmacol

10. Hundal RS et al. (2003) Oxidized low density lipoprotein inhibits macrophage apoptosis by blocking ceramide generation, thereby maintaining protein kinase B activation and Bcl-XL levels. J Biol Chem 278(27):24399–24408.

11. Getz GS, Reardon CA (2009) Apoprotein E as a lipid transport and signaling protein in the blood, liver, and artery wall. J Lipid Res 50 Suppl:S156–61.

12. Baitsch D et al. (2011) Apolipoprotein E induces antiinflammatory phenotype in macrophages. Arterioscler Thromb Vasc Biol 31(5):1160–1168.

13. Sung HY et al. (2012) Enhanced Macrophage Tribbles-1 Expression in Murine Experimental Atherosclerosis. Biology 1(1):43–57.

14. Varbo A, Benn M, Tybjaerg-Hansen A, Grande P, Nordestgaard BG (2011) TRIB1 and GCKR Polymorphisms, Lipid Levels, and Risk of Ischemic Heart Disease in the General Population. Arterioscler Thromb Vasc Biol 31(2):451–457.

15. Aulchenko YS et al. (2009) Loci influencing lipid levels and coronary heart disease risk in 16 European population cohorts. Nat Genet 41(1):47–55.

16. Bauer RC, Yenilmez BO, Rader DJ (2015) Tribbles-1: a novel regulator of hepatic lipid metabolism in humans. Biochem Soc Trans 43(5):1079–1084.

17. Satoh T et al. (2013) Critical role of Trib1 in differentiation of tissue-resident M2-like macrophages. Nature 495(7442):524–528.

18. Tao H et al. (2015) Macrophage SR-BI mediates efferocytosis via Src/PI3K/Rac1 signaling and reduces atherosclerotic lesion necrosis. J Lipid Res 56(8):1449–1460.

19. Clausen BE, Burkhardt C, Reith W, Renkawitz R, Förster I (1999) Conditional gene targeting in macrophages and granulocytes using LysMcre mice. Transgenic Res 8(4):265–277.

20. Drechsler M, Megens RT, van Zandvoort M, Weber C, Soehnlein O (2010) Hyperlipidemia-triggered neutrophilia promotes early atherosclerosis. Circulation 122(18):1837–1845.

21. Hasty AH et al. (1999) Retroviral gene therapy in ApoE-deficient mice: ApoE expression in the artery wall reduces early foam cell lesion formation. Circulation 99(19):2571–2576.

22. Bjørklund MM et al. (2014) Induction of atherosclerosis in mice and hamsters without germline genetic engineering. Circ Res 114(11):1684–1689.

23. Norata GD, Tavori H, Pirillo A, Fazio S, Catapano AL (2016) Biology of proprotein convertase subtilisin kexin 9: beyond low-density lipoprotein cholesterol lowering. Cardiovasc Res 112(1):429–442.

24. Arndt L et al. (2018) Tribbles homolog 1 deficiency modulates function and polarization of murine bone marrow-derived macrophages. J Biol Chem 293(29):11527–11536.

25. Schunkert H et al. (2011) Large-scale association analysis identifies 13 new susceptibility loci for coronary artery disease. Nat Genet 43(4):333–338.

26. Ference BA et al. (2017) Low-density lipoproteins cause atherosclerotic cardiovascular disease. 1. Evidence from genetic, epidemiologic, and clinical studies. A consensus statement from the European Atherosclerosis Society Consensus Panel. Eur Heart J 38(32):2459–2472.

27. Chui PC, Guan HP, Lehrke M, Lazar MA (2005) PPARgamma regulates adipocyte cholesterol metabolism via oxidized LDL receptor 1. J Clin Invest 115(8):2244–2256.

28. Schaeffer DF et al. (2009) LOX-1 augments oxLDL uptake by lysoPC-stimulated murine macrophages but is not required for oxLDL clearance from plasma. J Lipid Res 50(8):1676–1684.

29. Guan H et al. (2016) Competition between members of the tribbles pseudokinase protein family shapes their interactions with mitogen activated protein kinase pathways. Sci Rep 6:32667.

30. Hunerdosse DM et al. (2014) Chemical genetics screening reveals KIAA1363 as a cytokine-lowering target. ACS Chem Biol 9(12):2905–2913.

31. Rahman K et al. (2017) Inflammatory Ly6Chi monocytes and their conversion to M2 macrophages drive atherosclerosis regression. J Clin Invest 127(8):2904–2915.

32. Peled M et al. (2017) A wild-type mouse-based model for the regression of inflammation in atherosclerosis. PLoS One 12(3):e0173975.

33. Sasaki Y et al. (2006) Canonical NF-kappaB activity, dispensable for B cell development, replaces BAFF-receptor signals and promotes B cell proliferation upon activation. Immunity 24(6):729–739.

34. Houthuys E, Movahedi K, De Baetselier P, Van Ginderachter JA, Brouckaert P (2010) A method for the isolation and purification of mouse peripheral blood monocytes. J Immunol Methods 359(1-2):1–10.

35. Ghosn EE et al. (2010) Two physically, functionally, and developmentally distinct peritoneal macrophage subsets. Proc Natl Acad Sci U S A 107(6):2568–2573.

36. Stary HC et al. (1995) A definition of advanced types of atherosclerotic lesions and a histological classification of atherosclerosis. A report from the Committee on Vascular Lesions of the Council on Arteriosclerosis, American Heart Association. Arterioscler Thromb Vasc Biol 15(9):1512–1531.

37. Park YM (2014) CD36, a scavenger receptor implicated in atherosclerosis. Exp Mol Med 46:e99.

38. Chen Y et al. (2006) The di-leucine motif contributes to class a scavenger receptor-mediated internalization of acetylated lipoproteins. Arterioscler Thromb Vasc Biol 26(6):1317–1322.

39. Xue J et al. (2014) Transcriptome-based network analysis reveals a spectrum model of human macrophage activation. Immunity 40(2):274–288.

