## Supplementary figures and tables for "Myeloid Tribbles 1 induces early atherosclerosis via enhanced foam cell expansion"

### Supplemental Figure 1

**A**

#### "Knockout (KO)- first" *Trib1* null allele on C57BL/6 background

| | +/+ | +/tm1a | tm1a/tm1a |
|---|---|---|---|
| Observed | 14 | 32 | 0.0 |
| Expected | 11.5 | 23 | 11.5 |
| Chi-squared test | 0.0004 | | |

**B**

#### "Conditional-ready" Trib1 null allele

| | +/+ | +/tm1c | tm1c/tm1c |
|---|---|---|---|
| Observed | 9 | 17 | 10 |
| Expected | 9 | 18 | 9 |
| Chi-squared test | 0.92 | | |

**C**

#### Conditional-*Trib1* null allele (tm1d) on C57BL/6 background

| | tm1c/tm1c | tm1c/tm1d | tm1d/tm1d |
|---|---|---|---|
| Observed | 20 | 54 | 7 |
| Expected | 25.25 | 50.5 | 25.25 |
| Chi-squared test | 0.0007 | | |

**D**

#### "KO- first" *Trib1* null allele on mixed genetic background

| | +/+ | +/tm1a | tm1a/tm1a |
|---|---|---|---|
| Observed | 39 | 77 | 25 |
| Expected | 35.25 | 71 | 35.25 |
| Chi-squared test | 0.14 | | |

**E**

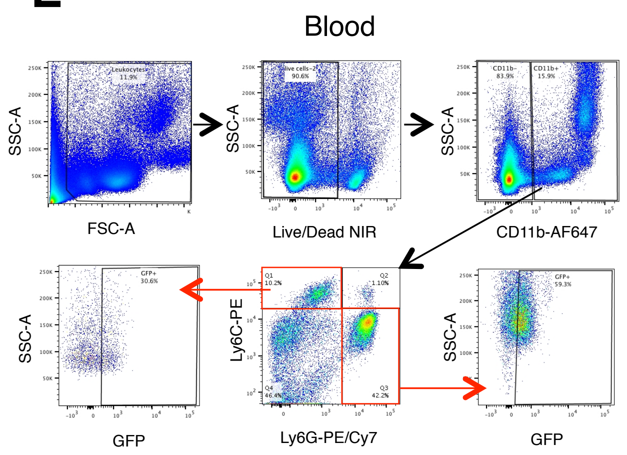

**F**

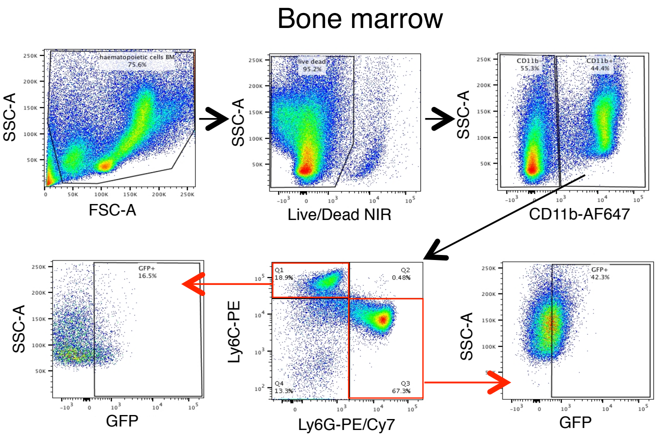

**G**

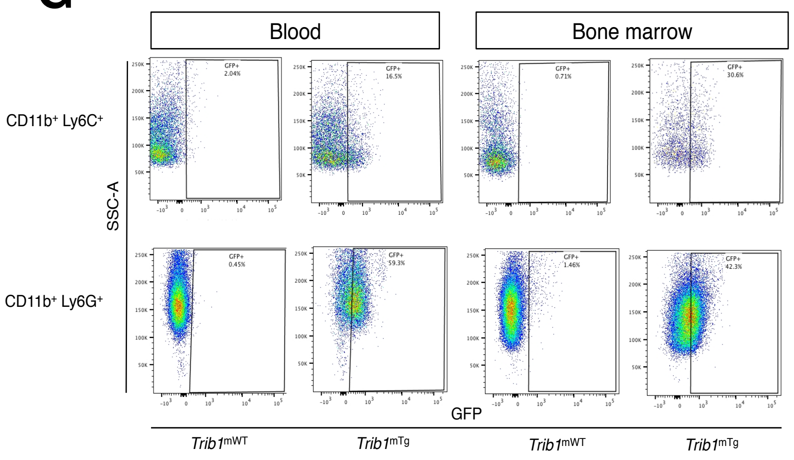

**H**

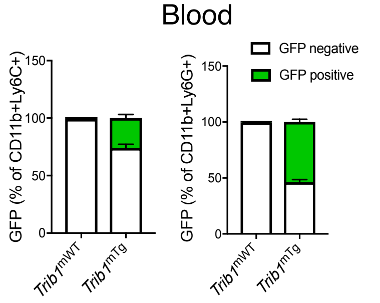

**I**

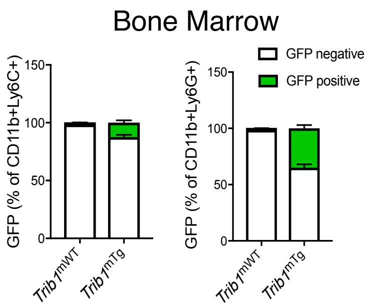

### Supplemental Figure 2

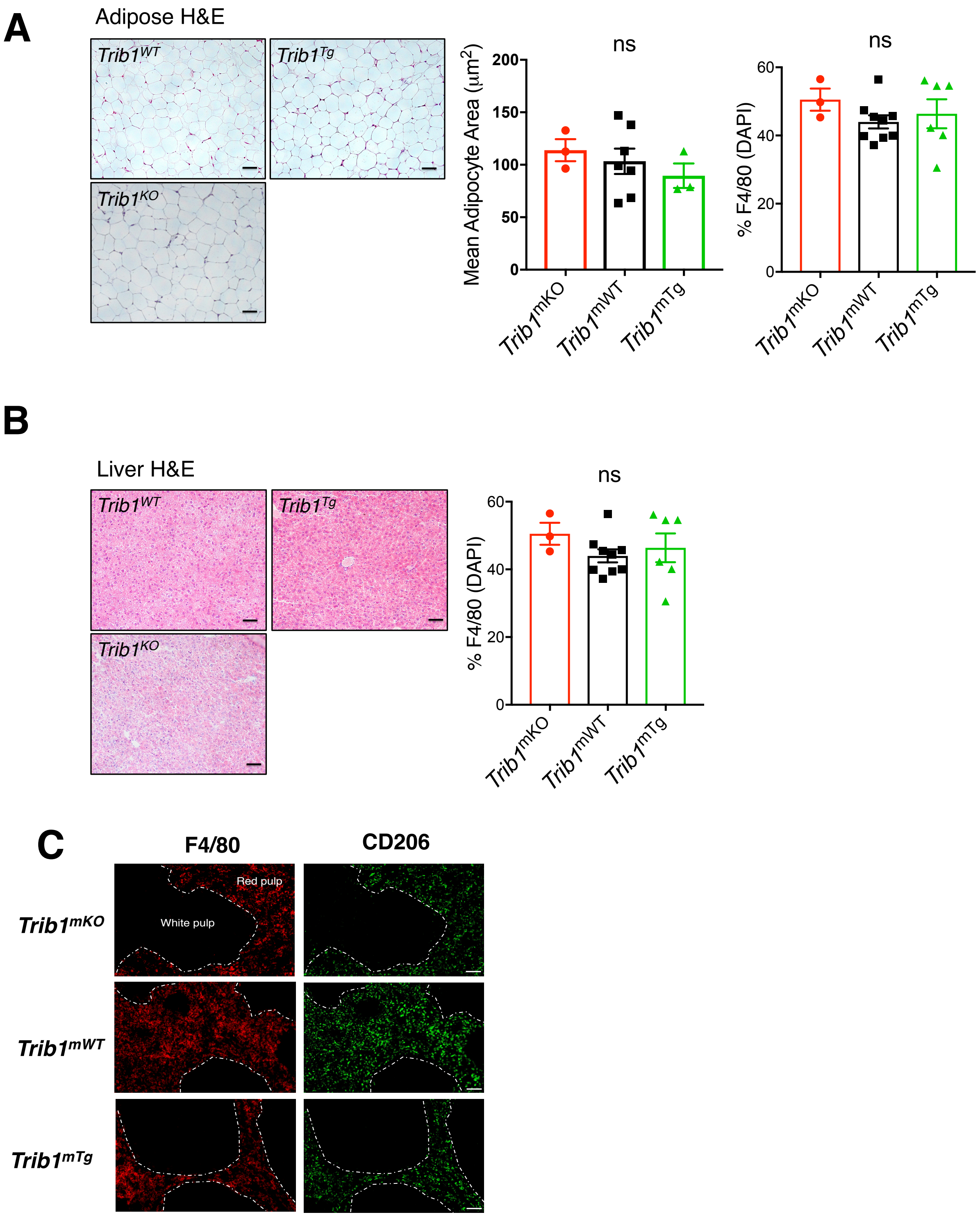

### Supplemental Figure 3

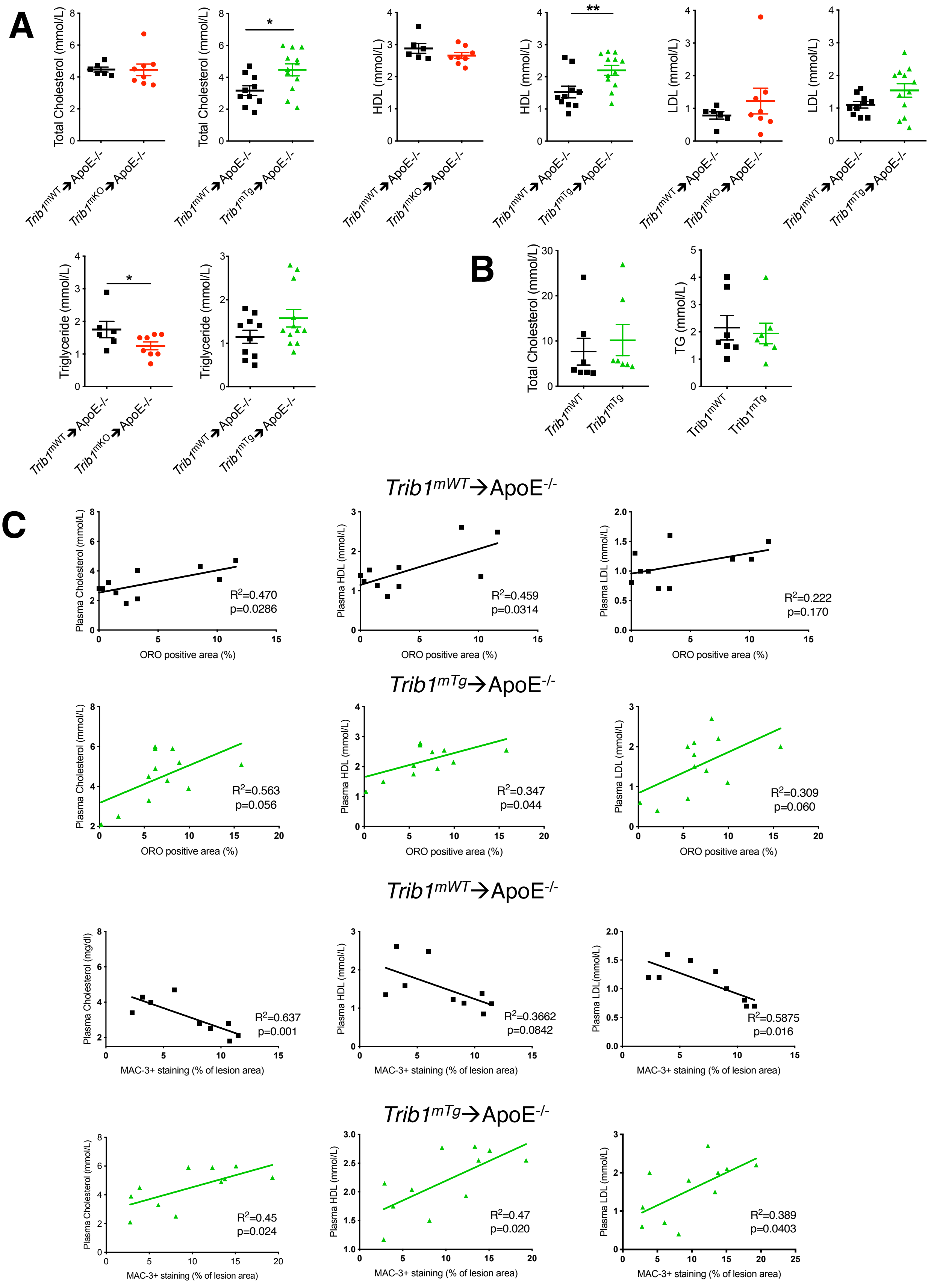

### Supplemental Figure 4

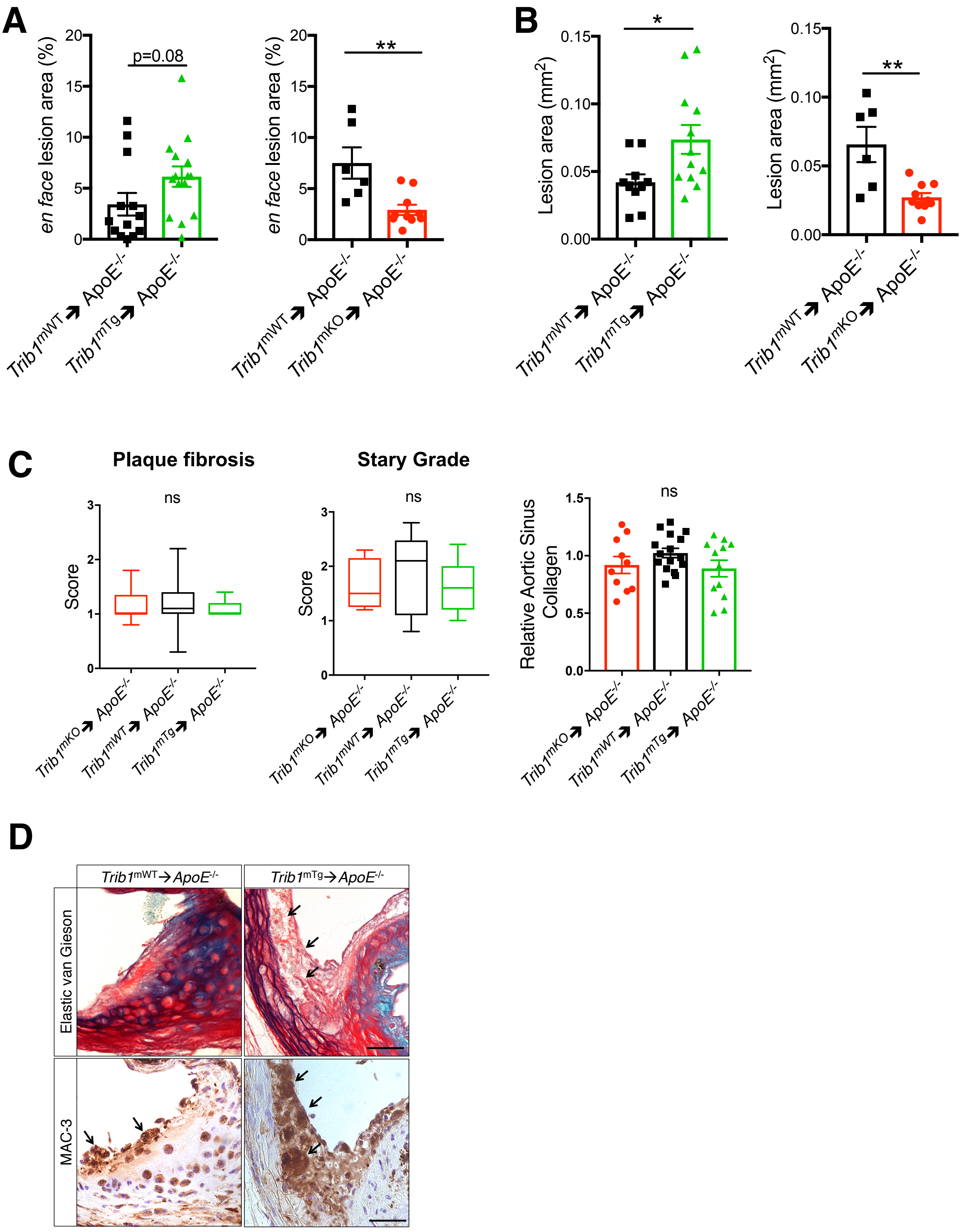

### Supplemental Figure 5

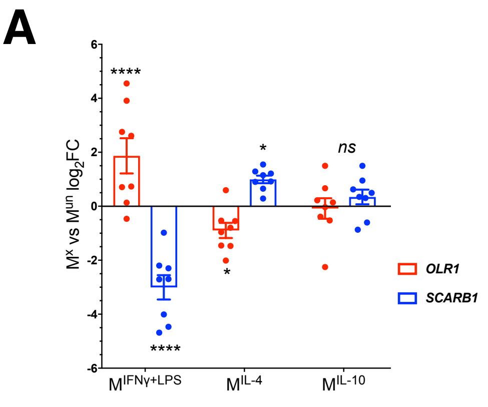

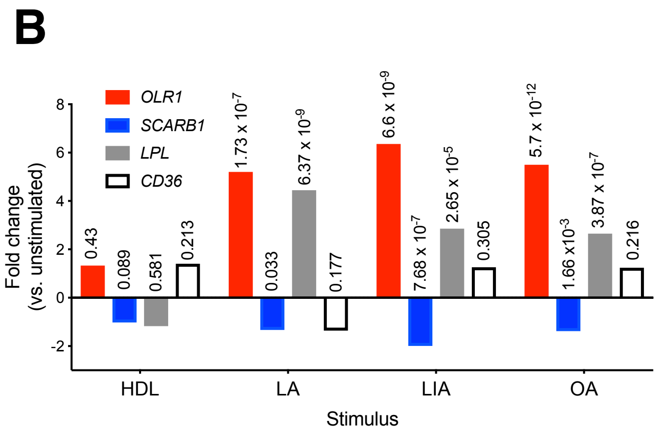

**STable. 1.**

| | Monocyte-derived-Macrophages | | | | | Monocytes | | | | |
|---|---|---|---|---|---|---|---|---|---|---|
| **Gene.ID** | **log.fold** | **fold.change** | **t.statistics** | **p.values** | **q.values** | **log.fold** | **fold.change** | **t.statistics** | **p.values** | **q.values** |
| AARS | -0.304137224 | 0.809926432 | -8.782897728 | 1.31E-16 | 2.69E-14 | -0.065066552 | 0.95590122 | -3.843905469 | 0.000142206 | 0.001565015 |
| ABHD14A | -0.220750797 | 0.858118744 | -8.037885032 | 2.20E-14 | 2.57E-12 | -0.098909216 | 0.9337387 | -5.156138259 | 4.09E-07 | 1.30E-05 |
| ACAT1 | -0.148057487 | 0.902464766 | -5.76394764 | 2.06E-08 | 5.23E-07 | -0.07291358 | 0.950716049 | -4.575812807 | 6.46E-06 | 0.000125567 |
| ADCY3 | -0.254850629 | 0.838073901 | -8.218052384 | 6.54E-15 | 9.18E-13 | -0.158691965 | 0.895836923 | -3.566095551 | 0.000409236 | 0.003567832 |
| **ADORA2B[B]** | **-0.38349133** | **0.766580215** | **-8.95060471** | **3.99E-17** | **8.90E-15** | **-0.102855836** | **0.931187866** | **-3.588055639** | **0.000377325** | **0.003367428** |
| AHR | 0.141652554 | 1.103168031 | 4.970874874 | 1.13E-06 | 1.79E-05 | 0.16153246 | 1.118474573 | 5.087764035 | 5.74E-07 | 1.68E-05 |
| AKR7A2 | -0.078918283 | 0.946767257 | -3.488746621 | 0.000559022 | 0.003634005 | -0.072663646 | 0.950880766 | -4.49269597 | 9.38E-06 | 0.00017144 |
| ALDH3A2 | -0.108851881 | 0.927325749 | -3.553768781 | 0.00044182 | 0.002983349 | -0.083274197 | 0.943913003 | -4.0872901 | 5.34E-05 | 0.000715008 |
| ALG3 | -0.085239414 | 0.942628095 | -4.042024821 | 6.76E-05 | 0.000611276 | -0.052242293 | 0.964436199 | -3.536274173 | 0.000456637 | 0.003915552 |
| ALKBH7 | -0.101966495 | 0.931762068 | -4.386983346 | 1.60E-05 | 0.000180532 | -0.077062159 | 0.94798612 | -4.923005676 | 1.28E-06 | 3.35E-05 |
| ANAPC11 | -0.085872707 | 0.942214405 | -4.44761784 | 1.23E-05 | 0.000141461 | -0.05873906 | 0.960102899 | -3.585411076 | 0.00038104 | 0.003378556 |
| ANKRD37 | 0.17271609 | 1.127178572 | 4.329594849 | 2.05E-05 | 0.000222413 | 0.107501134 | 1.077360543 | 4.524879725 | 8.12E-06 | 0.000152942 |
| ANO7 | 0.093401452 | 1.066882616 | 4.001031406 | 7.98E-05 | 0.000706169 | 0.101983961 | 1.073248355 | 4.397662424 | 1.43E-05 | 0.000245118 |
| ANTXR2 | 0.177347745 | 1.130803101 | 6.150376873 | 2.50E-09 | 8.29E-08 | 0.122042637 | 1.088274604 | 5.997331838 | 4.72E-09 | 2.77E-07 |
| ARHGAP19 | 0.128541933 | 1.093188306 | 5.633869829 | 4.10E-08 | 9.67E-07 | 0.060538818 | 1.042855174 | 3.79454163 | 0.000172403 | 0.001818891 |
| ARID4A | 0.078770827 | 1.056117847 | 3.729865936 | 0.000229583 | 0.001728025 | 0.062753195 | 1.04445707 | 3.724469238 | 0.000225804 | 0.00224404 |
| ARNTL | 0.077498448 | 1.055186819 | 4.150248844 | 4.35E-05 | 0.000424794 | 0.105821071 | 1.076106654 | 5.110912101 | 5.12E-07 | 1.54E-05 |
| ATF3 | 0.192376677 | 1.142644544 | 3.71259996 | 0.000245077 | 0.001822696 | 0.402352716 | 1.321661491 | 9.92633625 | 9.00E-21 | 2.30E-18 |
| ATP6V1F | -0.089599337 | 0.939783708 | -5.656464306 | 3.64E-08 | 8.79E-07 | -0.089599337 | 0.961284253 | -3.463365081 | 0.000595061 | 0.00485921 |
| ATPAF1 | -0.073672061 | 0.950216352 | -3.99988055 | 8.01E-05 | 0.00070889 | -0.064855686 | 0.956040946 | -4.070990188 | 5.71E-05 | 0.000758148 |
| BAZ1A | 0.074830962 | 1.053237623 | 3.207162652 | 0.001487371 | 0.008220651 | 0.093386614 | 1.066871643 | 4.589957134 | 6.06E-06 | 0.000119581 |
| **BCL3** | **0.239372378** | **1.180479001** | **6.103909899** | **3.24E-09** | **1.04E-07** | **0.099144323** | **1.071137971** | **3.292867123** | **0.001085954** | **0.007829639** |

| | | | | | | | | | | |
|---|---|---|---|---|---|---|---|---|---|---|
| BSCL2 | -0.19840344 | 0.871514492 | -5.868162062 | 1.18E-08 | 3.20E-07 | -0.107471522 | 0.928213431 | -3.947405377 | 9.43E-05 | 0.00113083 |
| BTG2 | 0.317979002 | 1.246583051 | 10.29838022 | 1.85E-21 | 1.16E-18 | 0.580118403 | 1.494971937 | 15.99920551 | 2.74E-44 | 2.14E-41 |
| C10orf59 | -0.158281039 | 0.896092123 | -5.486518069 | 8.81E-08 | 1.89E-06 | -0.072477235 | 0.951003638 | -3.68994754 | 0.000257517 | 0.002476517 |
| C12orf10 | -0.086294318 | 0.941939093 | -4.979681707 | 1.08E-06 | 1.74E-05 | -0.07205218 | 0.951283869 | -4.289748921 | 2.28E-05 | 0.000360624 |
| C12orf52 | -0.086294318 | 0.941939093 | -4.979681707 | 1.08E-06 | 1.74E-05 | -0.07205218 | 0.951283869 | -4.289748921 | 2.28E-05 | 0.000360624 |
| C13orf18 | 0.440171348 | 1.35676546 | 8.548328088 | 6.77E-16 | 1.16E-13 | 0.187277971 | 1.13861339 | 4.8072306 | 2.22E-06 | 5.18E-05 |
| C16orf7 | 0.174663087 | 1.128700788 | 8.090312857 | 1.55E-14 | 1.90E-12 | 0.09825619 | 1.070478774 | 4.426983893 | 1.25E-05 | 0.000219117 |
| C17orf101 | -0.130454337 | 0.913543709 | -5.174329622 | 4.23E-07 | 7.47E-06 | -0.057615887 | 0.960850654 | -3.309233856 | 0.001026123 | 0.00748459 |
| C3orf62 | 0.129900536 | 1.094218259 | 5.19385354 | 3.84E-07 | 6.90E-06 | 0.088406916 | 1.063195509 | 4.597797245 | 5.85E-06 | 0.00011629 |
| C5orf5 | 0.157784854 | 1.115572947 | 6.439594797 | 4.83E-10 | 1.92E-08 | 0.083170164 | 1.059343276 | 5.789485938 | 1.49E-08 | 7.56E-07 |
| C5orf54 | -0.112721816 | 0.924841591 | -4.320203558 | 2.13E-05 | 0.000229811 | -0.089628436 | 0.939764753 | -4.20260421 | 3.30E-05 | 0.000485327 |
| C7orf43 | 0.161297182 | 1.118292185 | 6.292869891 | 1.12E-09 | 4.00E-08 | 0.101608485 | 1.072969067 | 5.209504439 | 3.13E-07 | 1.02E-05 |
| C7orf50 | -0.235472322 | 0.849406868 | -6.936902928 | 2.53E-11 | 1.48E-09 | -0.145898378 | 0.903816389 | -6.128250632 | 2.25E-09 | 1.44E-07 |
| C8orf55 | -0.11191407 | 0.925359543 | -3.556928637 | 0.000436758 | 0.002956334 | -0.092692092 | 0.937771219 | -4.065064582 | 5.85E-05 | 0.000772729 |
| CALN1 | -0.080819265 | 0.94552056 | -4.500936477 | 9.74E-06 | 0.000115383 | -0.054749069 | 0.962761884 | -3.434654998 | 0.000659637 | 0.005242707 |
| CAPG | -0.16007424 | 0.894979015 | -5.987681627 | 6.15E-09 | 1.82E-07 | -0.087814056 | 0.940947375 | -3.291216034 | 0.001092166 | 0.007865358 |
| **CCL3** | **0.38427235** | **1.305201319** | **7.108286426** | **8.83E-12** | **5.78E-10** | **1.350292888** | **2.549638815** | **18.66363469** | **1.89E-55** | **3.94E-52** |
| **CCL3L1** | **0.396937255** | **1.316709655** | **6.695411776** | **1.08E-10** | **5.17E-09** | **1.183835398** | **2.271799317** | **16.59021024** | **9.55E-47** | **8.53E-44** |
| **CCL3L3** | **0.365258445** | **1.288112362** | **6.653247048** | **1.39E-10** | **6.42E-09** | **1.356130877** | **2.559977051** | **18.18283748** | **2.01E-53** | **3.58E-50** |
| **CCL4L1** | **0.457586749** | **1.373242821** | **4.221349367** | **3.23E-05** | **0.000328203** | **0.611835011** | **1.528201743** | **8.981062111** | **1.32E-17** | **2.47E-15** |
| **CCL8** | **0.443291864** | **1.359703288** | **4.679551789** | **4.38E-06** | **5.80E-05** | **0.30255706** | **1.233328452** | **5.075349313** | **6.10E-07** | **1.76E-05** |
| CCNG1 | -0.149122761 | 0.901798639 | -6.072737251 | 3.85E-09 | 1.20E-07 | -0.089554517 | 0.939812905 | -4.530583694 | 7.92E-06 | 0.00014953 |
| **CD44** | **0.139235411** | **1.101321291** | **3.642123264** | **0.000319136** | **0.002274711** | **0.201566466** | **1.149946279** | **6.311093458** | **7.82E-10** | **5.53E-08** |
| CDK4 | -0.199216252 | 0.871023621 | -12.63759225 | 1.43E-29 | 2.99E-26 | -0.070434918 | 0.952350857 | -4.043999928 | 6.38E-05 | 0.000827016 |
| CEBPD | 0.305118352 | 1.235519989 | 6.297963502 | 1.09E-09 | 3.92E-08 | 0.100837452 | 1.072395784 | 4.908522132 | 1.37E-06 | 3.56E-05 |
| CHD4 | 0.101034254 | 1.072542082 | 6.14094502 | 2.64E-09 | 8.65E-08 | 0.050802633 | 1.035841046 | 3.737161807 | 0.000215099 | 0.002161709 |
| CHMP4A | -0.085559725 | 0.942418833 | -4.873877062 | 1.79E-06 | 2.65E-05 | -0.085559725 | 0.942418833 | -4.873877062 | 1.79E-06 | 2.65E-05 |
| CHSY1 | 0.103633518 | 1.074476193 | 5.740719898 | 2.33E-08 | 5.89E-07 | 0.103633518 | 1.074476193 | 5.740719898 | 2.33E-08 | 5.89E-07 |
| CKLF | 0.074151743 | 1.052741877 | 3.477736813 | 0.000581543 | 0.003753458 | 0.050100111 | 1.035336765 | 3.260562118 | 0.001213668 | 0.008541134 |
| COMMD1 | -0.082609651 | 0.944347896 | -4.604258254 | 6.15E-06 | 7.78E-05 | -0.047869247 | 0.967363999 | -3.71226477 | 0.000236569 | 0.002332486 |
| COPE | -0.075498487 | 0.949014157 | -4.5646801 | 7.34E-06 | 9.05E-05 | -0.065378476 | 0.955694568 | -4.605004898 | 5.66E-06 | 0.000113709 |

| | | | | | | | | | | |
|---|---|---|---|---|---|---|---|---|---|---|
| CSNK1D | 0.108421214 | 1.078047849 | 4.826411032 | 2.23E-06 | 3.22E-05 | 0.065883588 | 1.046725817 | 3.490105454 | 0.000540274 | 0.004500007 |
| CUTA | -0.108198671 | 0.927745709 | -5.822249075 | 1.51E-08 | 3.98E-07 | -0.057230143 | 0.961107598 | -3.528101273 | 0.000470499 | 0.004017879 |
| **CXCL1** | **0.208085468** | **1.155154216** | **3.718390564** | **0.000239774** | **0.001786445** | **0.198932598** | **1.147848786** | **5.388836063** | **1.26E-07** | **4.79E-06** |
| CYB5R1 | -0.098718605 | 0.933862075 | -3.406825538 | 0.000748195 | 0.004646763 | -0.093376272 | 0.937326599 | -4.823836772 | 2.05E-06 | 4.87E-05 |
| DDB1 | -0.125380561 | 0.916762183 | -6.437434951 | 4.89E-10 | 1.94E-08 | -0.059148032 | 0.95983077 | -4.002319239 | 7.56E-05 | 0.000937202 |
| DDOST | -0.101472284 | 0.932081308 | -6.219947453 | 1.69E-09 | 5.86E-08 | -0.057537999 | 0.960902529 | -3.678439906 | 0.000268988 | 0.00256318 |
| DKFZP586I1420 | 0.127170188 | 1.092149374 | 3.999501173 | 8.02E-05 | 0.000708965 | 0.061911947 | 1.043848215 | 3.211191243 | 0.001435988 | 0.009815595 |
| DLG4 | 0.185386394 | 1.137121485 | 5.193134341 | 3.85E-07 | 6.91E-06 | 0.125289074 | 1.090726261 | 5.017737027 | 8.09E-07 | 2.25E-05 |
| DPH5 | -0.080067353 | 0.946013481 | -3.677591437 | 0.000279567 | 0.002037985 | -0.106135083 | 0.929073679 | -5.056742829 | 6.69E-07 | 1.91E-05 |
| DPP7 | -0.18572505 | 0.879207105 | -5.217748514 | 3.41E-07 | 6.20E-06 | -0.086494053 | 0.941808695 | -3.248800407 | 0.001263528 | 0.008824934 |
| DUSP2 | 0.251213063 | 1.190207458 | 4.527844672 | 8.65E-06 | 0.000104142 | 0.267930924 | 1.204079729 | 8.267138396 | 2.41E-15 | 3.83E-13 |
| DUSP5 | 0.345577117 | 1.270659179 | 5.615939962 | 4.50E-08 | 1.05E-06 | 0.248670417 | 1.18811165 | 6.113623253 | 2.45E-09 | 1.55E-07 |
| DUSP6 | 0.617173415 | 1.533867023 | 10.84644877 | 2.66E-23 | 2.56E-20 | 0.127606397 | 1.092479643 | 4.585820364 | 6.17E-06 | 0.000121206 |
| DYRK4 | -0.083253511 | 0.943926537 | -3.277634972 | 0.00117149 | 0.006764882 | -0.065051267 | 0.955911348 | -3.570305093 | 0.000402929 | 0.003525139 |
| ECE2 | -0.096879762 | 0.935053124 | -3.192732554 | 0.001561088 | 0.008518869 | -0.082940526 | 0.94413134 | -3.565808922 | 0.000409669 | 0.003569116 |
| EGR1 | 0.599880182 | 1.51559069 | 9.430304225 | 1.24E-18 | 4.43E-16 | 1.258557259 | 2.392563572 | 19.80372323 | 2.96E-60 | 9.25E-57 |
| EGR2 | 0.202074918 | 1.150351628 | 4.861143979 | 1.90E-06 | 2.79E-05 | 1.419755997 | 2.675402581 | 20.29344534 | 2.56E-62 | 1.07E-58 |
| EI24 | -0.079329175 | 0.946497648 | -4.519105676 | 8.99E-06 | 0.000107518 | -0.051303562 | 0.965063941 | -3.457970981 | 0.000606724 | 0.004919106 |
| ENOPH1 | -0.074301387 | 0.949801943 | -4.475165863 | 1.09E-05 | 0.000127401 | -0.059563646 | 0.9595543 | -4.272964366 | 2.45E-05 | 0.000380763 |
| EPRS | -0.15026855 | 0.901082715 | -5.932946066 | 8.30E-09 | 2.39E-07 | -0.061128713 | 0.958513919 | -4.263150673 | 2.55E-05 | 0.000393711 |
| ESYT1 | -0.190315618 | 0.876413968 | -9.160527393 | 8.84E-18 | 2.63E-15 | -0.072486285 | 0.950997672 | -4.392510335 | 1.46E-05 | 0.000250022 |
| ETFB | -0.072510777 | 0.950981528 | | 0.001029572 | 0.00608304 | -0.093150543 | 0.937473267 | | 1.81E-07 | 6.53E-06 |
| EXOSC8 | -0.083520291 | 0.943752005 | -3.929664884 | 0.000105948 | 0.000896631 | -0.081954207 | 0.944777029 | -4.589360716 | 6.07E-06 | 0.000119581 |
| FAHD1 | -0.07766145 | 0.947592411 | -3.285440488 | 0.001140629 | 0.006635713 | -0.095970654 | 0.93564253 | -4.476082801 | 1.01E-05 | 0.000183254 |
| FAM100B | 0.212890808 | 1.159008228 | 7.558018214 | 5.16E-13 | 4.48E-11 | 0.137711156 | 1.100158323 | 5.632013658 | 3.50E-08 | 1.60E-06 |
| FAM118B | -0.039101255 | 0.973261064 | -2.099472163 | 0.036621846 | 0.103678967 | -0.074048542 | 0.949968419 | -3.367241862 | 0.000837856 | 0.00638323 |
| FAM195A | -0.100127511 | 0.93295053 | -3.657724447 | 0.000301123 | 0.002167319 | -0.105636622 | 0.929394736 | -4.066454887 | 5.82E-05 | 0.000770772 |
| FBXO6 | -0.13334427 | 0.911715577 | -5.566207073 | 5.84E-08 | 1.33E-06 | -0.082624987 | 0.944337858 | -3.373696546 | 0.000819025 | 0.006262664 |
| FBXW4 | -0.086993261 | 0.941482863 | -4.159134352 | 4.19E-05 | 0.000412415 | -0.06923469 | 0.95314348 | -3.66070969 | 0.000287609 | 0.002703522 |
| FCGR2A | 0.32179637 | 1.249885874 | 5.160308831 | 4.53E-07 | 7.90E-06 | 0.117075212 | 1.084533947 | 4.202750242 | 3.30E-05 | 0.000485327 |
| FGD4 | 0.114919685 | 1.082914758 | 3.898180416 | 0.000119935 | 0.000995637 | 0.155052231 | 1.113461929 | 8.36898061 | 1.16E-15 | 1.94E-13 |

| | | | | | | | | | | |
|---|---|---|---|---|---|---|---|---|---|---|
| **FOS** | **0.607372682** | **1.52348224** | **12.69986367** | **8.57E-30** | **2.14E-26** | **0.673150675** | **1.59455148** | **18.78229875** | **5.99E-56** | **1.50E-52** |
| FOSB | 0.139458541 | 1.101491637 | 3.227001728 | 0.001391276 | 0.007765888 | 1.072955623 | 2.103738847 | 16.79284667 | 1.36E-47 | 1.31E-44 |
| **FPR2** | **0.241812782** | **1.18247754** | **3.634792169** | **0.000327947** | **0.002325577** | **0.137361934** | **1.099892049** | **3.936734577** | **9.85E-05** | **0.001174557** |
| FTSJ2 | -0.076121105 | 0.948604682 | -3.340020497 | 0.000945058 | 0.005664003 | -0.083432251 | 0.943809598 | -4.214313827 | 3.14E-05 | 0.000464933 |
| FUCA1 | -0.210440731 | 0.864273163 | -3.672468488 | 0.000284982 | 0.002072625 | -0.139875517 | 0.907597464 | -4.562902241 | 6.85E-06 | 0.000131676 |
| FUT4 | 0.083736683 | 1.059759341 | 4.226694855 | 3.16E-05 | 0.000322511 | 0.115221443 | 1.083141287 | 5.858180876 | 1.02E-08 | 5.43E-07 |
| GABARAPL1 | 0.116479255 | 1.084086034 | 5.081546127 | 6.64E-07 | 1.12E-05 | 0.209678591 | 1.156430522 | 7.69540838 | 1.26E-13 | 1.64E-11 |
| GALIG | -0.286607221 | 0.819827778 | -6.488811919 | 3.63E-10 | 1.50E-08 | -0.079678483 | 0.946268507 | -3.329186033 | 0.000957329 | 0.007077784 |
| GALK1 | -0.12262555 | 0.918514531 | -6.161118128 | 2.35E-09 | 7.89E-08 | -0.044274495 | 0.969777377 | -3.210798353 | 0.0014379 | 0.00982329 |
| GNPDA1 | -0.080019859 | 0.946044624 | -3.310231335 | 0.00104755 | 0.006177582 | -0.07734392 | 0.947800995 | -4.305833639 | 2.13E-05 | 0.000340155 |
| GSTP1 | -0.146655732 | 0.903342047 | -6.930011316 | 2.63E-11 | 1.53E-09 | -0.076741426 | 0.948196895 | -4.838159566 | 1.92E-06 | 4.62E-05 |
| GTPBP6 | -0.095082032 | 0.936219012 | -4.005697416 | 7.83E-05 | 0.000697101 | -0.09003803 | 0.939497983 | -4.428348747 | 1.25E-05 | 0.000218109 |
| H3F3B | 0.272530861 | 1.207924984 | 9.179393049 | 7.71E-18 | 2.41E-15 | 0.231557338 | 1.174101667 | 13.2209369 | 5.06E-33 | 2.34E-30 |
| **HBEGF** | **0.76361341** | **1.697737499** | **10.26440275** | **2.40E-21** | **1.43E-18** | **0.546514932** | **1.460553224** | **12.21082218** | **4.22E-29** | **1.51E-26** |
| HDDC3 | -0.08217459 | 0.944632718 | -3.739321844 | 0.000221494 | 0.001677239 | -0.082304525 | 0.944547644 | -4.780695413 | 2.51E-06 | 5.72E-05 |
| HDHD2 | -0.090442368 | 0.939234711 | -4.118348867 | 4.96E-05 | 0.000471872 | -0.071304686 | 0.951776879 | -3.717504091 | 0.00023189 | 0.00229904 |
| HEATR1 | -0.073536338 | 0.950305749 | -3.514536055 | 0.000509426 | 0.00335379 | -0.107852143 | 0.927968576 | -4.336499811 | 1.86E-05 | 0.000307164 |
| HEBP1 | -0.240535578 | 0.84643103 | -8.917098822 | 5.06E-17 | 1.09E-14 | -0.066761639 | 0.954778749 | -4.30180895 | 2.16E-05 | 0.000345477 |
| HECA | 0.107416644 | 1.07729745 | 3.952383836 | 9.68E-05 | 0.000831447 | 0.067760897 | 1.048088757 | 3.574215195 | 0.000397153 | 0.003484356 |
| HMGN2 | -0.078167397 | 0.947260153 | -3.284264147 | 0.001145231 | 0.00664702 | -0.063817873 | 0.956728929 | -3.248606943 | 0.001264364 | 0.008825841 |
| IARS | -0.159480841 | 0.895347207 | -3.828475122 | 0.000157377 | 0.001248428 | -0.031125924 | 0.978656226 | -2.011458458 | 0.044992061 | 0.135409424 |
| IDH1 | -0.140574103 | 0.907158091 | -6.634017257 | 1.55E-10 | 7.08E-09 | -0.057706131 | 0.960790551 | -3.782111794 | 0.00018091 | 0.001888821 |
| IER2 | 0.13043528 | 1.094623913 | 5.14783564 | 4.81E-07 | 8.38E-06 | 0.549308043 | 1.463383647 | 14.72511368 | 4.73E-39 | 2.82E-36 |
| IGFBP6 | 0.369741009 | 1.29212085 | 6.313516874 | 9.96E-10 | 3.62E-08 | 0.084196228 | 1.060096963 | 3.815459833 | 0.000158934 | 0.001707035 |
| IIP45 | 0.10876382 | 1.078303891 | 3.18207532 | 0.001617687 | 0.008793184 | 0.088757659 | 1.063454021 | 3.533027491 | 0.000462097 | 0.003959656 |
| **IL1R2** | **0.535136879** | **1.449079633** | **6.47478239** | **3.94E-10** | **1.61E-08** | **0.371667432** | **1.293847364** | **5.229013227** | **2.84E-07** | **9.49E-06** |
| **IL1RN** | **0.439358621** | **1.356001356** | **5.922585133** | **8.78E-09** | **2.51E-07** | **0.080060824** | **1.057062605** | **3.58684789** | **0.000379017** | **0.003367428** |
| IMP4 | -0.105262596 | 0.929635717 | -4.660880202 | 4.77E-06 | 6.22E-05 | -0.061355292 | 0.958363394 | -3.423122372 | 0.000687373 | 0.005428638 |
| INSIG1 | 0.193337718 | 1.143405962 | 3.713048242 | 0.000244662 | 0.001821781 | 0.169977104 | 1.12504063 | 4.859926899 | 1.73E-06 | 4.25E-05 |
| INTS3 | 0.103182485 | 1.07414033 | 5.160299818 | 4.53E-07 | 7.90E-06 | 0.086771678 | 1.061991102 | 5.234630776 | 2.76E-07 | 9.27E-06 |
| IQSEC1 | 0.12637329 | 1.091546273 | 5.633176387 | 4.12E-08 | 9.69E-07 | 0.062226696 | 1.044075974 | 3.914810181 | 0.00010746 | 0.001260318 |

| | | | | | | | | | | |
|---|---|---|---|---|---|---|---|---|---|---|
| ITPRIP | 0.150137503 | 1.10967523 | 5.616852915 | 4.48E-08 | 1.05E-06 | 0.12930644 | 1.093767758 | 6.8158206 | 3.75E-11 | 3.76E-09 |
| JUNB | 0.303513913 | 1.234146716 | 8.533676014 | 7.49E-16 | 1.27E-13 | 0.528021491 | 1.441950351 | 14.33023008 | 1.89E-37 | 1.03E-34 |
| JUND | 0.091852088 | 1.065737466 | 4.325519724 | 2.08E-05 | 0.000225438 | 0.190486606 | 1.141148549 | 10.46591196 | 1.16E-22 | 3.31E-20 |
| **KDM6B** | **0.134598218** | **1.097787044** | **3.779738623** | **0.000189856** | **0.001467891** | **0.135263623** | **1.098293486** | **4.643552159** | **4.74E-06** | **9.87E-05** |
| KIAA0247 | 0.150819664 | 1.11020005 | 6.861205738 | 4.00E-11 | 2.21E-09 | 0.067099528 | 1.047608396 | 3.713980573 | 0.000235027 | 0.002322776 |
| KLK7 | 0.097379743 | 1.069828649 | 4.375426099 | 1.68E-05 | 0.000187718 | 0.109934104 | 1.079178943 | 5.125490873 | 4.76E-07 | 1.46E-05 |
| LCTL | -0.073382755 | 0.95040692 | -3.524884704 | 0.000490709 | 0.003251231 | -0.077897206 | 0.947437574 | -3.830133355 | 0.000150085 | 0.00162456 |
| LGALS3 | -0.302035982 | 0.811106925 | -8.011439364 | 2.63E-14 | 3.01E-12 | -0.095920296 | 0.935675189 | -3.766314934 | 0.000192294 | 0.001973778 |
| LHPP | -0.187064511 | 0.87839119 | -4.410614108 | 1.44E-05 | 0.000164483 | -0.123066078 | 0.918234104 | -5.12094427 | 4.87E-07 | 1.48E-05 |
| **LILRB2** | **0.281246918** | **1.215244764** | **5.874882157** | **1.14E-08** | **3.10E-07** | **0.07711268** | **1.054904706** | **4.273221879** | **2.45E-05** | **0.000380763** |
| LRRC41 | -0.111440257 | 0.925663501 | -3.729580825 | 0.000229831 | 0.001728852 | -0.084673072 | 0.942998204 | -3.586684787 | 0.000379246 | 0.003367428 |
| LRRC42 | -0.081547692 | 0.945043281 | -4.421582938 | 1.38E-05 | 0.000157415 | -0.052551539 | 0.964229491 | -3.385452136 | 0.000785739 | 0.006048832 |
| LRRC8C | 0.15506471 | 1.11347156 | 4.456437389 | 1.18E-05 | 0.000136621 | 0.148366148 | 1.108313596 | 5.092897854 | 5.60E-07 | 1.66E-05 |
| LSMD1 | -0.08660218 | 0.941738111 | -4.270120473 | 2.63E-05 | 0.000274272 | -0.065416821 | 0.955669167 | -3.590226823 | 0.0003743 | 0.003349675 |
| **LYN** | **0.124278318** | **1.089962363** | **5.19724828** | **3.78E-07** | **6.81E-06** | **0.110701754** | **1.079753321** | **6.094186571** | **2.73E-09** | **1.72E-07** |
| MACROD1 | -0.156854275 | 0.896978758 | -4.67686575 | 4.43E-06 | 5.84E-05 | -0.07415812 | 0.949896268 | -3.334758741 | 0.000938896 | 0.006982795 |
| MAFB | 0.144881451 | 1.105639797 | 3.313932327 | 0.001034277 | 0.006107949 | 0.21383436 | 1.159766491 | 8.72348876 | 8.92E-17 | 1.56E-14 |
| MAP1LC3A | 0.428650969 | 1.345974399 | 8.976695553 | 3.31E-17 | 7.80E-15 | 0.193058166 | 1.143184425 | 4.721697731 | 3.31E-06 | 7.22E-05 |
| MAP3K7IP2 | 0.086358869 | 1.06168727 | 3.27720179 | 0.001173225 | 0.006771774 | 0.083928212 | 1.059900042 | 3.738332282 | 0.000214136 | 0.002159982 |
| MAPKAPK2 | 0.088045712 | 1.062929353 | 3.230745927 | 0.001373798 | 0.007690295 | 0.096529561 | 1.069198383 | 4.088519704 | 5.31E-05 | 0.000712153 |
| MCL1 | 0.113188597 | 1.081616149 | 4.497360535 | 9.89E-06 | 0.000116992 | 0.242797132 | 1.18328462 | 10.53243128 | 6.75E-23 | 2.01E-20 |
| MCTP2 | 0.160146787 | 1.117400822 | 4.228528075 | 3.14E-05 | 0.000320567 | 0.15025669 | 1.109766908 | 3.724563015 | 0.000225723 | 0.00224404 |
| MFNG | -0.154402077 | 0.89850468 | -6.075223264 | 3.80E-09 | 1.19E-07 | -0.063995929 | 0.956610857 | -4.649530344 | 4.62E-06 | 9.64E-05 |
| MGAM | 0.098685744 | 1.07079755 | 3.32953704 | 0.000980012 | 0.005845474 | 0.184650242 | 1.136541404 | 4.182544336 | 3.59E-05 | 0.000518822 |
| MIDN | 0.125505973 | 1.090890256 | 5.564850505 | 5.88E-08 | 1.33E-06 | 0.156433573 | 1.114528549 | 8.266422367 | 2.42E-15 | 3.83E-13 |
| MRPL48 | -0.104044308 | 0.930421082 | -3.66205612 | 0.000296293 | 0.002141191 | -0.076004498 | 0.948681358 | -4.005017281 | 7.48E-05 | 0.000929398 |
| MSL3 | 0.097200767 | 1.069695937 | 3.910750445 | 0.000114153 | 0.000952697 | 0.062734517 | 1.044443548 | 3.926668085 | 0.000102499 | 0.00121349 |
| MUSK | -0.094047769 | 0.936890425 | -4.882702196 | 1.71E-06 | 2.57E-05 | -0.061788393 | 0.958075733 | -3.416592166 | 0.000703557 | 0.005528514 |
| MXD1 | 0.122354214 | 1.088509662 | 3.19508589 | 0.00154884 | 0.008470515 | 0.238570051 | 1.179822682 | 6.647594105 | 1.05E-10 | 9.27E-09 |
| MYST3 | 0.10262168 | 1.073722871 | 3.367407341 | 0.000859143 | 0.005229313 | 0.133614279 | 1.097038593 | 5.215322173 | 3.04E-07 | 1.01E-05 |
| NDUFAF2 | -0.106635181 | 0.928751679 | -4.894592217 | 1.62E-06 | 2.44E-05 | -0.065102849 | 0.955877171 | -3.695443793 | 0.000252201 | 0.002446978 |

| | | | | | | | | | | |
|---|---|---|---|---|---|---|---|---|---|---|
| NDUFS3 | -0.110578468 | 0.926216608 | -4.935707217 | 1.34E-06 | 2.07E-05 | -0.114339926 | 0.923804881 | -8.126974477 | 6.46E-15 | 9.85E-13 |
| NENF | -0.156982628 | 0.896898959 | -5.421725918 | 1.23E-07 | 2.52E-06 | -0.074490091 | 0.949677717 | -3.234726814 | 0.001325682 | 0.00916686 |
| **NFKBIZ** | **0.347874164** | **1.272683923** | **7.267250879** | **3.28E-12** | **2.38E-10** | **0.45296965** | **1.368855016** | **17.01467971** | **1.61E-48** | **1.83E-45** |
| **NFX1** | **-0.089432833** | **0.939892177** | **-5.033396996** | **8.38E-07** | **1.38E-05** | **-0.056539121** | **0.961568059** | **-3.622581058** | **0.000331836** | **0.003048213** |
| NIT2 | -0.101571536 | 0.932017187 | -4.748863094 | 3.19E-06 | 4.42E-05 | -0.086244685 | 0.941971499 | -4.765017102 | 2.70E-06 | 6.07E-05 |
| **NLRP3** | **0.221092397** | **1.165615847** | **4.100077147** | **5.34E-05** | **0.000502789** | **0.076727127** | **1.054622826** | **4.343716761** | **1.81E-05** | **0.000300174** |
| NMRAL1 | -0.085638949 | 0.942367083 | -3.353984697 | 0.000900298 | 0.005442712 | -0.080977831 | 0.945416644 | -4.640768734 | 4.81E-06 | 9.96E-05 |
| NPTN | 0.099185878 | 1.071168824 | 6.70178539 | 1.04E-10 | 5.00E-09 | 0.043258145 | 1.030438316 | 3.398397344 | 0.000750544 | 0.005817299 |
| NR2C2AP | -0.100623338 | 0.932629948 | -3.917973783 | 0.000110951 | 0.000929187 | -0.088165754 | 0.940718021 | -4.161258603 | 3.93E-05 | 0.000558092 |
| NSMCE2 | -0.152545494 | 0.899661697 | -6.588525854 | 2.03E-10 | 8.90E-09 | -0.059179036 | 0.959810143 | -4.029542009 | 6.77E-05 | 0.000864092 |
| NUBP1 | -0.085481669 | 0.942469823 | -4.02009873 | 7.39E-05 | 0.000663469 | -0.052435844 | 0.96430682 | -4.081918351 | 5.46E-05 | 0.000730235 |
| **OSM** | **0.321101239** | **1.24928379** | **5.800203377** | **1.70E-08** | **4.42E-07** | **0.700639974** | **1.625225576** | **15.24872599** | **3.42E-41** | **2.25E-38** |
| PDCL3 | -0.082740008 | 0.944262572 | -3.824633135 | 0.000159733 | 0.001265513 | -0.050043586 | 0.965907147 | -3.267262309 | 0.001186087 | 0.008401392 |
| PDE4B | 0.327127017 | 1.254512649 | 5.714525947 | 2.68E-08 | 6.72E-07 | 0.160562852 | 1.117723121 | 4.065153201 | 5.85E-05 | 0.000772729 |
| PDIA5 | -0.193318073 | 0.874591921 | -5.466876833 | 9.74E-08 | 2.06E-06 | -0.095870481 | 0.935707498 | -4.398492832 | 1.42E-05 | 0.000244593 |
| PDLIM4 | 0.178257976 | 1.131516777 | 3.871595217 | 0.000133091 | 0.001088233 | 0.068099831 | 1.048335015 | 3.27579928 | 0.001151787 | 0.008233069 |
| PDS5B | 0.079997806 | 1.057016433 | 3.381533147 | 0.000817731 | 0.005013863 | 0.06686481 | 1.047437971 | 3.764787918 | 0.00019343 | 0.001983805 |
| PDXP | -0.118804266 | 0.920950636 | -5.028452585 | 8.58E-07 | 1.41E-05 | -0.089541741 | 0.939821228 | -4.275920652 | 2.42E-05 | 0.000377386 |
| PEMT | -0.137906939 | 0.908836739 | -5.568717429 | 5.76E-08 | 1.32E-06 | -0.068638674 | 0.953537331 | -3.740529268 | 0.00021234 | 0.002149535 |
| PEPD | -0.130715773 | 0.913378178 | -6.822591139 | 5.05E-11 | 2.71E-09 | -0.048453321 | 0.966972442 | -3.315473055 | 0.001004132 | 0.007345616 |
| PHACTR2 | 0.114677963 | 1.082733332 | 4.647679352 | 5.06E-06 | 6.57E-05 | 0.096806678 | 1.069403778 | 4.486815565 | 9.63E-06 | 0.0001754 |
| PIK3CD | 0.075858421 | 1.053987985 | 3.304209788 | 0.001069486 | 0.006295061 | 0.063236491 | 1.044807017 | 3.651845782 | 0.000297366 | 0.002776453 |
| PIK3CG | 0.128019611 | 1.092792593 | 3.948743786 | 9.82E-05 | 0.000840657 | 0.102561997 | 1.073678453 | 3.896477026 | 0.000115579 | 0.001336713 |
| PLCB2 | 0.14419825 | 1.105116335 | 4.633572923 | 5.39E-06 | 6.94E-05 | 0.11826077 | 1.085425547 | 5.143517986 | 4.35E-07 | 1.36E-05 |
| **PLEK** | **0.183221455** | **1.135416376** | **5.253313602** | **2.86E-07** | **5.31E-06** | **0.083788268** | **1.059797234** | **3.292007573** | **0.001089184** | **0.007848401** |
| PLEKHG2 | 0.162183288 | 1.118979254 | 4.789346908 | 2.65E-06 | 3.75E-05 | 0.088080544 | 1.062955016 | 3.909007539 | 0.00010997 | 0.001282742 |
| PMAIP1 | 0.156305945 | 1.114429957 | 4.084295661 | 5.70E-05 | 0.000529856 | 0.340874612 | 1.266524172 | 8.756108303 | 7.02E-17 | 1.25E-14 |
| PNRC1 | 0.111342396 | 1.080232902 | 4.014727395 | 7.55E-05 | 0.000675048 | 0.189766999 | 1.140579493 | 8.085652817 | 8.63E-15 | 1.28E-12 |
| POFUT2 | -0.081974139 | 0.944763976 | -4.229119756 | 3.13E-05 | 0.000320035 | -0.057976816 | 0.960610301 | -3.34313745 | 0.000911801 | 0.006834137 |
| POLR1C | -0.074944427 | 0.94937869 | -3.623946854 | 0.000341401 | 0.002407333 | -0.094174239 | 0.936808299 | -5.554022368 | 5.30E-08 | 2.26E-06 |
| PPA2 | -0.07420225 | 0.949867213 | -3.287428391 | 0.001132891 | 0.006596835 | -0.082983799 | 0.944103021 | -5.08803853 | 5.73E-07 | 1.68E-05 |

| | | | | | | | | | | |
|---|---|---|---|---|---|---|---|---|---|---|
| PPM1B | 0.123715347 | 1.089537118 | 3.907006002 | 0.000115847 | 0.000964904 | 0.197387686 | 1.146620268 | 6.837722535 | 3.28E-11 | 3.33E-09 |
| PPP1R14A | 0.101993702 | 1.073255601 | 4.513812499 | 9.20E-06 | 0.000109745 | 0.096762875 | 1.069371309 | 3.737777955 | 0.000214592 | 0.002160668 |
| PRDX4 | -0.122274476 | 0.918738075 | -5.354852024 | 1.72E-07 | 3.40E-06 | -0.063036505 | 0.957247236 | -3.935635899 | 9.89E-05 | 0.001178603 |
| PRKCA | 0.168047696 | 1.123537047 | 4.118823258 | 4.95E-05 | 0.000471313 | 0.117445353 | 1.084812234 | 3.731002385 | 0.000220233 | 0.00219916 |
| PRPF38B | 0.074819111 | 1.053228971 | 3.887352861 | 0.000125137 | 0.001029932 | 0.082492927 | 1.05884611 | 4.870438567 | 1.64E-06 | 4.10E-05 |
| PSCD4 | 0.16455209 | 1.120818046 | 6.458496931 | 4.33E-10 | 1.75E-08 | 0.085713665 | 1.061212567 | 3.780685422 | 0.000181912 | 0.001890489 |
| PSMG1 | -0.163159796 | 0.893066929 | -6.982837056 | 1.91E-11 | 1.15E-09 | -0.055318577 | 0.962381906 | -3.34189826 | 0.000915762 | 0.006855604 |
| **PTGER4** | **0.168539459** | **1.123920086** | **4.900698401** | **1.58E-06** | **2.38E-05** | **0.126426806** | **1.091586764** | **5.544644821** | **5.57E-08** | **2.34E-06** |
| PTGES2 | -0.079042844 | 0.946685517 | -3.526549762 | 0.000487758 | 0.003241867 | -0.129356795 | 0.91423896 | -7.036804222 | 9.41E-12 | 1.01E-09 |
| PTGS2 | 0.141803996 | 1.103283838 | 3.516518304 | 0.00050579 | 0.003333361 | 0.885610294 | 1.847546011 | 18.06600062 | 6.22E-53 | 9.72E-50 |
| PTPN1 | 0.076072969 | 1.054144738 | 4.045351584 | 6.67E-05 | 0.000604506 | 0.05852511 | 1.041400576 | 3.959030897 | 9.00E-05 | 0.001088681 |
| RARA | 0.204796892 | 1.152524078 | 5.642775463 | 3.91E-08 | 9.30E-07 | 0.080018282 | 1.057031435 | 4.019030828 | 7.06E-05 | 0.000892163 |
| RFX2 | 0.122455843 | 1.088586344 | 3.959551171 | 9.41E-05 | 0.000812012 | 0.225426628 | 1.169122926 | 6.070595993 | 3.12E-09 | 1.94E-07 |
| RGL2 | 0.130432094 | 1.094621497 | 5.221786219 | 3.34E-07 | 6.10E-06 | 0.122475235 | 1.088600976 | 5.925625387 | 7.05E-09 | 3.86E-07 |
| RGS2 | 0.229580862 | 1.172494261 | 4.696722507 | 4.05E-06 | 5.45E-05 | 0.168739653 | 1.124076056 | 8.334394873 | 1.49E-15 | 2.42E-13 |
| RNF113A | -0.082422209 | 0.944470598 | -3.258936807 | 0.001248599 | 0.007095446 | -0.116217893 | 0.92260314 | -5.092753527 | 5.60E-07 | 1.66E-05 |
| RNF149 | 0.096919046 | 1.069487074 | 3.731752536 | 0.000227947 | 0.001717781 | 0.131518445 | 1.095446058 | 4.738500744 | 3.06E-06 | 6.76E-05 |
| RNF217 | 0.127029555 | 1.092042917 | 5.112944546 | 5.70E-07 | 9.74E-06 | 0.084240061 | 1.060129172 | 3.81238199 | 0.000160851 | 0.001718771 |
| RNPEP | -0.177302645 | 0.884354898 | -4.931097549 | 1.37E-06 | 2.10E-05 | -0.11881042 | 0.920946708 | -3.65914325 | 0.000289311 | 0.002715439 |
| RNU2-1 | 0.089136211 | 1.0637331 | 3.154657464 | 0.001772137 | 0.009435798 | 0.232229619 | 1.174648913 | 4.8201538 | 2.09E-06 | 4.92E-05 |
| RPP21 | -0.094224611 | 0.93677559 | -4.38143606 | 1.64E-05 | 0.000183906 | -0.056630747 | 0.961506991 | -3.770796441 | 0.000188998 | 0.001951156 |
| RPUSD3 | -0.090199345 | 0.939392939 | -5.022377427 | 8.84E-07 | 1.45E-05 | -0.057699648 | 0.960794869 | -4.009102015 | 7.36E-05 | 0.000916105 |
| S100A6 | -0.099892632 | 0.933102432 | -5.116743895 | 5.60E-07 | 9.60E-06 | -0.053921892 | 0.963314047 | -3.269435636 | 0.001177266 | 0.008376881 |
| SARS2 | -0.106524186 | 0.928823136 | -4.087708421 | 5.62E-05 | 0.000523705 | -0.092005007 | 0.93821794 | -3.362818804 | 0.000850992 | 0.006467541 |
| SEC11C | -0.119082005 | 0.920773358 | -4.287702423 | 2.45E-05 | 0.000257837 | -0.100547919 | 0.932678703 | -4.750360269 | 2.90E-06 | 6.43E-05 |
| SERPINB6 | -0.165039723 | 0.891903962 | -6.598549199 | 1.91E-10 | 8.45E-09 | -0.072064957 | 0.951275444 | -4.208462677 | 3.22E-05 | 0.000476011 |
| SERTAD1 | 0.084364397 | 1.060220541 | 3.98495583 | 8.51E-05 | 0.000743052 | 0.074447725 | 1.052957878 | 4.179484188 | 3.64E-05 | 0.000524785 |
| SH2B1 | 0.09856572 | 1.07070847 | 3.932551086 | 0.000104746 | 0.000889027 | 0.078067408 | 1.055603038 | 3.435970305 | 0.000656541 | 0.005225323 |
| SLC18A1 | 0.098355691 | 1.070552606 | 3.791154669 | 0.000181724 | 0.001418174 | 0.0990115 | 1.071039361 | 3.338810689 | 0.000925701 | 0.006905097 |
| SLC22A4 | 0.319186567 | 1.247626902 | 6.6158757 | 1.73E-10 | 7.74E-09 | 0.096331762 | 1.069051802 | 3.837675718 | 0.00014572 | 0.0015897 |
| SLC25A25 | 0.091011409 | 1.065116626 | 3.779264106 | 0.000190201 | 0.001469652 | 0.100590778 | 1.072212439 | 4.019269151 | 7.06E-05 | 0.000892163 |

| | | | | | | | | | | |
|---|---|---|---|---|---|---|---|---|---|---|
| SLC25A37 | 0.192104326 | 1.142428857 | 4.991330908 | 1.03E-06 | 1.66E-05 | 0.171914461 | 1.126552432 | 4.10071437 | 5.05E-05 | 0.000685251 |
| SLC39A11 | -0.129511434 | 0.91414097 | -4.785507913 | 2.70E-06 | 3.80E-05 | -0.071631819 | 0.951561087 | -3.913258903 | 0.000108125 | 0.001264532 |
| SLC47A1 | -0.438124158 | 0.73809368 | -7.513785862 | 6.85E-13 | 5.79E-11 | -0.153173451 | 0.899270189 | -3.913505762 | 0.000108019 | 0.001264473 |
| SLC7A1 | -0.215612978 | 0.861180181 | -5.23197469 | 3.18E-07 | 5.86E-06 | -0.088604519 | 0.940431965 | -4.202197849 | 3.31E-05 | 0.000485327 |
| SMU1 | -0.091258964 | 0.938703235 | -3.685538694 | 0.000271358 | 0.00198809 | -0.093463147 | 0.937270157 | -3.996267862 | 7.75E-05 | 0.000954403 |
| SNF8 | -0.109213554 | 0.927093304 | -3.561845788 | 0.000428989 | 0.002913213 | -0.090036648 | 0.939498883 | -3.594467712 | 0.000368457 | 0.003316378 |
| SNTA1 | -0.101696995 | 0.93193614 | -4.957948309 | 1.20E-06 | 1.89E-05 | -0.079569109 | 0.946340249 | -4.281244365 | 2.36E-05 | 0.000371664 |
| SNTB1 | -0.211121373 | 0.863865508 | -6.269608528 | 1.28E-09 | 4.51E-08 | -0.07764309 | 0.947604471 | -3.247651786 | 0.001268498 | 0.008845789 |
| SON | 0.075373826 | 1.053634015 | 3.141570888 | 0.001850561 | 0.009790825 | 0.254330085 | 1.192781747 | 11.63066244 | 6.57E-27 | 2.16E-24 |
| SORL1 | 0.217433749 | 1.162663612 | 4.243383738 | 2.95E-05 | 0.000302432 | 0.155869881 | 1.114093164 | 5.317985172 | 1.81E-07 | 6.53E-06 |
| **SP100** | **0.074333565** | **1.052874561** | **3.147790262** | **0.001812899** | **0.009628234** | **0.10708263** | **1.077048062** | **4.86147341** | **1.72E-06** | **4.23E-05** |
| SPATA2L | 0.077883117 | 1.055468203 | 3.306623063 | 0.001060644 | 0.006248902 | 0.082802203 | 1.059073123 | 3.657107 | 0.000291538 | 0.00273019 |
| SRGN | 0.120913519 | 1.087423204 | 3.566045338 | 0.000422457 | 0.002875101 | 0.091791169 | 1.065692465 | 5.033841392 | 7.48E-07 | 2.12E-05 |
| SSH1 | 0.117886582 | 1.08514406 | 3.275325526 | 0.001180769 | 0.006801375 | 0.092150307 | 1.065957787 | 3.528921079 | 0.000469091 | 0.004011335 |
| SSR4 | -0.154674864 | 0.898334805 | -7.311714215 | 2.48E-12 | 1.87E-10 | -0.081743484 | 0.944915035 | -5.285984039 | 2.13E-07 | 7.53E-06 |
| STRA13 | -0.102395792 | 0.931484849 | -4.550799609 | 7.81E-06 | 9.54E-05 | -0.068303416 | 0.953758943 | -4.469907997 | 1.04E-05 | 0.000186722 |
| STX8 | -0.106395069 | 0.928906267 | -4.97611805 | 1.10E-06 | 1.76E-05 | -0.088080028 | 0.94077392 | -4.768060974 | 2.67E-06 | 6.01E-05 |
| TCEB2 | -0.12807382 | 0.915052346 | -7.164054602 | 6.25E-12 | 4.24E-10 | -0.053410397 | 0.963655642 | -3.897313069 | 0.000115197 | 0.001335985 |
| TGFA | 0.207188734 | 1.154436432 | 4.902585253 | 1.56E-06 | 2.36E-05 | 0.25810278 | 1.195904992 | 7.682987411 | 1.37E-13 | 1.77E-11 |
| THOC6 | -0.102747106 | 0.931258048 | -4.935452565 | 1.34E-06 | 2.07E-05 | -0.050463218 | 0.965626238 | -3.313130403 | 0.001012337 | 0.007396983 |
| TLE4 | 0.099582303 | 1.071463202 | 3.962745287 | 9.29E-05 | 0.000803427 | 0.095692412 | 1.068578142 | 6.151186144 | 1.97E-09 | 1.28E-07 |
| **TLR1** | **0.212698796** | **1.158853983** | **5.920077987** | **8.90E-09** | **2.54E-07** | **0.144246725** | **1.105153468** | **5.979914611** | **5.21E-09** | **3.01E-07** |
| **TLR6** | **0.118636299** | **1.085708117** | **5.635337649** | **4.07E-08** | **9.62E-07** | **0.191448304** | **1.141909489** | **9.762034106** | **3.30E-20** | **8.08E-18** |
| TMED3 | -0.156932298 | 0.89693025 | -6.041643793 | 4.57E-09 | 1.40E-07 | -0.077397534 | 0.947765773 | -3.694631609 | 0.00025298 | 0.002449853 |
| TMEM108 | -0.083008478 | 0.944086871 | -3.226132034 | 0.001395365 | 0.007784405 | -0.075176839 | 0.949225762 | -3.406084436 | 0.000730346 | 0.00570318 |
| TMEM119 | 0.407320849 | 1.326220673 | 4.852267483 | 1.98E-06 | 2.90E-05 | 0.090267444 | 1.064567511 | 3.323504339 | 0.000976469 | 0.007198008 |
| TMEM126B | -0.09773921 | 0.934496257 | -4.190760703 | 3.67E-05 | 0.000366626 | -0.073910422 | 0.950059371 | -3.814685801 | 0.000159414 | 0.001709726 |
| TMEM185B | 0.087540945 | 1.062557523 | 4.512213377 | 9.26E-06 | 0.000110417 | 0.109293468 | 1.078699835 | 5.545711283 | 5.54E-08 | 2.33E-06 |
| TMEM4 | -0.077282717 | 0.947841204 | -4.299799302 | 2.32E-05 | 0.000246188 | -0.048000547 | 0.967275963 | -3.318931382 | 0.000992132 | 0.007283404 |
| TMEM43 | 0.113626321 | 1.081944369 | 5.403639932 | 1.34E-07 | 2.74E-06 | 0.080609124 | 1.057464421 | 5.069355037 | 6.28E-07 | 1.80E-05 |
| TMEM86B | 0.132776113 | 1.096401428 | 4.832430923 | 2.17E-06 | 3.16E-05 | 0.099500867 | 1.071402722 | 5.01787758 | 8.08E-07 | 2.25E-05 |

| | | | | | | | | | | |
|---|---|---|---|---|---|---|---|---|---|---|
| TMEM88 | 0.113122368 | 1.081566497 | 3.774864035 | 0.000193432 | 0.001488176 | 0.210288183 | 1.156919259 | 7.519166485 | 4.11E-13 | 4.98E-11 |
| TMUB1 | 0.10526693 | 1.075693399 | 5.09535617 | 6.21E-07 | 1.06E-05 | 0.061799654 | 1.04376697 | 3.308876704 | 0.001027395 | 0.007485344 |
| TNFAIP3 | 0.278513353 | 1.212944343 | 8.39573452 | 1.94E-15 | 3.11E-13 | 0.131558543 | 1.095476505 | 4.412551789 | 1.34E-05 | 0.000231533 |
| TOMM40 | -0.156370358 | 0.897279678 | -4.358870061 | 1.81E-05 | 0.000199781 | -0.067701725 | 0.9541568 | -3.598877343 | 0.000362472 | 0.003281412 |
| TP53INP2 | 0.071232699 | 1.050613989 | 4.410001098 | 1.45E-05 | 0.000164771 | 0.087800336 | 1.062748583 | 4.915240647 | 1.33E-06 | 3.47E-05 |
| TRAPPC2L | -0.102562753 | 0.931377055 | -3.813981748 | 0.000166441 | 0.001307883 | -0.099801477 | 0.933161391 | -5.347559655 | 1.55E-07 | 5.71E-06 |
| TRAPPC6A | -0.139423253 | 0.907882027 | -4.798217477 | 2.54E-06 | 3.61E-05 | -0.0762047 | 0.948549719 | -3.248868851 | 0.001263233 | 0.008824934 |
| TRIB3 | -0.505640758 | 0.70434748 | -7.257304883 | 3.49E-12 | 2.46E-10 | -0.097255745 | 0.93480947 | -3.551400268 | 0.000431986 | 0.003732334 |
| TSPAN4 | -0.221314221 | 0.857783684 | -4.710113765 | 3.81E-06 | 5.18E-05 | -0.158522667 | 0.895942054 | -5.253527858 | 2.51E-07 | 8.60E-06 |
| TXNRD2 | -0.150498275 | 0.900939244 | -5.915606515 | 9.12E-09 | 2.58E-07 | -0.067631696 | 0.954203117 | -3.461327303 | 0.000599442 | 0.004875878 |
| UBE2D4 | -0.146220347 | 0.903614705 | -6.845449803 | 4.40E-11 | 2.41E-09 | -0.066175723 | 0.955166588 | -3.609610983 | 0.000348285 | 0.003173654 |
| UBP1 | 0.091088361 | 1.06517344 | 4.827129867 | 2.22E-06 | 3.22E-05 | 0.081511559 | 1.058126093 | 4.307660668 | 2.11E-05 | 0.000339171 |
| UQCRC2 | -0.123570753 | 0.91791295 | -4.830210277 | 2.19E-06 | 3.18E-05 | -0.060680077 | 0.958812035 | -3.387712162 | 0.000779485 | 0.00601551 |
| UROD | -0.131555255 | 0.912846852 | -5.453197036 | 1.04E-07 | 2.19E-06 | -0.061655979 | 0.958163672 | -3.676231608 | 0.000271244 | 0.002572909 |
| **VARS** | **-0.085127761** | **0.94270105** | **-3.400475399** | **0.000765115** | **0.004728356** | **-0.084732456** | **0.942959389** | **-4.659160596** | **4.42E-06** | **9.25E-05** |
| VNN3 | 0.161419074 | 1.118386673 | 3.457440708 | 0.000625299 | 0.003982418 | 0.2947784 | 1.226696541 | 6.390723 | 4.90E-10 | 3.74E-08 |
| WASPIP | 0.104509289 | 1.075128639 | 4.490045346 | 1.02E-05 | 0.00012036 | 0.091601082 | 1.065552061 | 4.799488687 | 2.30E-06 | 5.35E-05 |
| WBP11 | 0.090806583 | 1.064965417 | 4.702803815 | 3.94E-06 | 5.33E-05 | 0.084585568 | 1.06038309 | 4.75980026 | 2.77E-06 | 6.20E-05 |
| YARS | -0.157195994 | 0.896766323 | -7.66595984 | 2.56E-13 | 2.36E-11 | -0.040546757 | 0.972286397 | -3.569282261 | 0.000404453 | 0.003533526 |
| YIF1A | -0.107681134 | 0.928078579 | -3.821679167 | 0.000161567 | 0.001276809 | -0.079763628 | 0.946212662 | -3.640561673 | 0.000310238 | 0.002884527 |
| YPEL5 | 0.075428218 | 1.053673739 | 3.846044248 | 0.000147015 | 0.001178947 | 0.103422692 | 1.074319188 | 6.630585064 | 1.17E-10 | 1.01E-08 |
| ZBTB24 | -0.084531568 | 0.9430907 | -4.770365716 | 2.89E-06 | 4.05E-05 | -0.097970295 | 0.934346585 | -6.759323447 | 5.32E-11 | 5.12E-09 |
| ZBTB3 | -0.104260947 | 0.930281378 | -4.153533406 | 4.29E-05 | 0.000420087 | -0.116240583 | 0.922588629 | -5.281923335 | 2.17E-07 | 7.67E-06 |
| ZEB2 | 0.10072671 | 1.072313469 | 3.421123906 | 0.000711368 | 0.004464619 | 0.15967798 | 1.11703778 | 6.970708709 | 1.43E-11 | 1.51E-09 |
| ZMYND15 | 0.253099048 | 1.191764392 | 6.232534497 | 1.58E-09 | 5.49E-08 | 0.131169235 | 1.095180933 | 3.740695801 | 0.000212204 | 0.002149535 |
| ZSWIM4 | 0.45012701 | 1.366160524 | 7.895154562 | 5.70E-14 | 5.99E-12 | 0.50124646 | 1.415435943 | 15.78706282 | 2.07E-43 | 1.52E-40 |

[A]Please see Methods for cut-off values for definition of differential expression. The differentially expressed MDM gene list contained 2171 genes compared to 1842 in the monocytes. Only those that display concordant differential expression *in* $TRIB1^{High}$ versus $TRIB1^{Low}$ cells are shown.

[B]Grey highlighting indicates those Genes assigned by DAVID to the GO 0006954 ‘Inflammatory Response’ cluster. There was a 2.84-fold over-representation of these genes in this ‘Cluster of Terms but P values (P= 6.96 x $10^{-5}$, Benjamini-Hochberg adjusted *P* value 1.058 X $10^{-1}$) were modest

**STable. 2. Top Ranking Biological Processes[A] Enriched in Differentially Expressed Gene Lists[B] of: (1) Human *TRIB1*$^{High}$ versus *TRIB*1$^{Low}$ monocytes and (2) between *TRIB1*$^{HIgh}$ versus *TRIB*1$^{Low}$ Monocyte Derived Macrophages (MDMs).**

| | Monocytes | | | | Monocyte Derived Macrophages | | | |
|---|---|---|---|---|---|---|---|---|
| **Cluster description[C]** | **Rank** | **Cluster Gene Ontology terms in differentially expressed gene-sets (DEG[C])** | **Cluster enrichment in DEG of *TRIB1*$^{High}$ monocytes** | **p value[D] ranges** | **Rank** | **Cluster Gene Ontology terms in differentially expressed gene-sets (DEG[C])** | **Cluster enrichment in DEG of *TRIB1*$^{High}$ MDMs** | **p value[D] ranges** |
| Ribonucleoprotein Complex | 1 | 0022613; 0042254; 0034660; 0016072; 0006364; 0034470 | 8.78 | $3.9 \times 10^{-9}$- $2.6 \times 10^{-5}$ | - | - | - | - |
| Oxoacid, Carboxylic & Fatty acid metabolism | - | | | | 1 | 0043436; 0019752; 0006631 | 8.28 | $7.9 \times 10^{-8}$ - $3.3 \times 10^{-3}$ |
| Apoptosis & Cell Death | 2 | 0006915[E]; 0012501; 0016265; 0008219 | 6.87 | $6.8 \times 10^{-6}$ - $1.6 \times 10^{-4}$ | - | | - | - |
| Signal transduction & Cell communication | - | | - | - | 2 | 0009966; 0010648; 0009968; 1902532 | 7.62 | $1.1 \times 10^{-10}$- $3.4 \times 10^{-4}$ |
| Regulation of Cell Death & Apoptosis | 3 | 00043067[E]; 0042981[E]; 0043069[E]; 0060548[E]; 0043066[E]; 0043068[E]; 0010942[E]; 0043065[E]; 0010941; 0006916; 0006917; 0012502 | 6.03 | $6.5 \times 10^{-7}$- $5.2 \times 10^{-3}$ | 4 | 0043067; 0042981; 0043069; 0006915; 0060548; 0043066; 0043068; 0010942; 0043065; | 6.81 | $8.4 \times 10^{-8}$- $1.48 \times 10^{-3}$ |
| Response to unfolded proteins | 13 | 0006986[F] | 2.53 | $8 \times 10^{-3}$ | 3 | 0006986; 0034620; 0030968; 0036498; 0035967 | 6.91 | $9.0 \times 10^{-8}$- $1.48 \times 10^{-4}$ |
| Macromolecular Complex Subunit Organisation | 4 | 0034621; 0034622; 0043933; 0065003; 0070271 | 5.72 | $7.33 \times 10^{-6}$ - $4.82 \times 10^{-2}$ | - | - | - | |

[A] Annotation was performed using the Database for Annotation, Visualization and Integrated Discovery (DAVID) Platform.

[B] DEG lists, defined as in the Methods Section, comprised 1842 (monocytes) and 2171 (MDM) genes.

[C] The monocyte and MDM DEG lists returned respectively 13 and 27 GO clusters with Benjamini-Hochberg adjusted P values > $5.0 \times 10^{-2}$

[D] Benjamini-Hochberg adjusted P values

[E] GO term also present in MDM Cluster 4 of GO terms

[F] GO term also present in MDM Cluster 3 of GO terms

**STable. 3** The most significantly altered pathways in *TRIB1*$^{HIGH}$ versus *TRIB1*$^{LOW}$ macrophages**.**

| Monocyte Derived Macrophages | | | | | | Monocytes | | |
|---|---|---|---|---|---|---|---|---|
| **Pathway No.** | **Pathway Name** | **Rank-ing** | **Log. fold change** | **p value** | **FDR** | **Log. fold change** | **p value** | **FDR** |
| 645 | Creation of C4 and C2 Activators | 1 | 0.34 | $1.6 \times 10^{-5}$ | $7.3 \times 10^{-5}$ | Not present | | |
| 199 | Translocation of Zap 70 to Immunological Synapse | 2 | 0.20 | $< 10^{-7}$ | $< 10^{-7}$ | -0.03 | 0.45 | 0.54 |
| 380 | PD1 Signaling | 3 | 0.17 | $< 10^{-7}$ | $< 10^{-7}$ | -0.02 | 0.60 | 0.67 |
| 197 | Phosphorylation Of CD3 And TCR Zeta Chains | 4 | 0.16 | $< 10^{-7}$ | $< 10^{-7}$ | -0.03 | 0.46 | 0.54 |
| 222 | **HDL Mediated Lipid Transport** | 5 | 0.16 | $< 10^{-7}$ | $< 10^{-7}$ | 0.01 | 0.36 | 0.43 |
| 263 | **Chemokine Receptors Bind Chemokines** | 6 | 0.15 | $< 10^{-7}$ | $< 10^{-7}$ | 0.23 | $< 10^{-7}$ | $< 10^{-7}$ |
| 647 | Initial Triggering of Complement | 7 | 0.15 | $5.1 \times 10^{-4}$ | $1.7 \times 10^{-3}$ | 0.01 | 0.74 | 0.79 |
| 202 | Generation of Second Messenger Molecules | 8 | 0.13 | $< 10^{-7}$ | $< 10^{-7}$ | -0.01 | 0.59 | 0.67 |
| 604 | **Lipoprotein Metabolism** | 9 | 0.11 | $< 10^{-7}$ | $< 10^{-7}$ | Not present | | |
| 270 | Norepinephrine Neurotransmitter Release Cycle | 10 | 0.10 | $1.9 \times 10^{-6}$ | $9.5 \times 10^{-6}$ | 0.04 | $1 \times 10^{-3}$ | $2 \times 10^{-3}$ |
| 280 | Steroid Hormones | 11 | 0.09 | $< 10^{-7}$ | $< 10^{-7}$ | -0.01 | 0.45 | 0.54 |
| 24 | Metabolism of Steroid Hormones and Vitamins A and D | 12 | 0.09 | $< 10^{-7}$ | $< 10^{-7}$ | -0.01 | 0.22 | 0.29 |
| 246 | **Peptide Ligand Binding Receptors** | 13 | 0.09 | $< 10^{-7}$ | $< 10^{-7}$ | 0.11 | $< 10^{-7}$ | $< 10^{-7}$ |
| 405 | Zinc Transporters | 14 | 0.08 | $2.3 \times 10^{-6}$ | $1.1 \times 10^{-5}$ | -0.02 | 0.03 | 0.05 |
| 607 | **Chylomicron Mediated Lipid Transport** | 15 | 0.08 | $6.2 \times 10^{-4}$ | $2.0 \times 10^{-3}$ | $-4.35 \times 10^{-3}$ | 0.61 | 0.68 |
| - - - | - - - | - - - | - - - | - - - | - - - | - - - | - - - | - - - |
| 244 | **Mitochondrial fatty acid Beta oxidation** | 10 | -0.07 | $1.8 \times 10^{-7}$ | $1.0 \times 10^{-6}$ | -0.04 | $1.2 \times 10^{-6}$ | $5.4 \times 10^{-6}$ |
| 278 | **tRNA aminoacylation** | 9 | -0.07 | $< 10^{-7}$ | $< 10^{-7}$ | -0.04 | $< 10^{-7}$ | $< 10^{-7}$ |
| 85 | Glycosphingolipid Metabolism | 8 | -0.07 | $< 10^{-7}$ | $< 10^{-7}$ | -0.01 | 0.17 | 0.23 |
| 313 | CDC6 association with the ORC origin complex | 7 | -0.08 | $3.3 \times 10^{-4}$ | $1.0 \times 10^{-3}$ | 0.02 | 0.27 | 0.34 |
| 221 | Apoptotic cleavage of cell adhesion proteins | 6 | -0.08 | $6.0 \times 10^{-6}$ | $2.8 \times 10^{-5}$ | 0.01 | 0.40 | 0.49 |
| 257 | **Cytosolic tRNA Aminoacylation** | 5 | -0.09 | $< 10^{-7}$ | $< 10^{-7}$ | -0.03 | $< 10^{-7}$ | $1.9 \times 10^{-7}$ |
| 661 | **Metabolism of Porphyrins** | 4 | -0.10 | $< 10^{-7}$ | $< 10^{-7}$ | -0.04 | $6.7 \times 10^{-5}$ | $2.1 \times 10^{-4}$ |
| 300 | Hormone ligand binding receptors | 3 | -0.10 | $1.5 \times 10^{-6}$ | $7.8 \times 10^{-6}$ | -0.01 | 0.33 | 0.41 |
| 558 | Ethanol oxidation | 2 | -0.13 | $< 10^{-7}$ | $< 10^{-7}$ | -0.01 | 0.46 | 0.55 |
| 485 | Amino acid synthesis and interconversion transamination | 1 | -0.17 | $< 10^{-7}$ | $< 10^{-7}$ | -0.02 | 0.07 | 0.11 |

Results of Quantitative Set Analysis of Gene Expression (Qusage version 2.2.2) analysis comparing gene expression values of genes assigned to the specified Reactome Pathways in *TRIB1*$^{High}$ versus *TRIB1*$^{Low}$ samples in CTS dataser.

Ranking of Reactome pathways (version 4.0) according to MDM transcriptome.

Increased (above dotted line) and reduced average RNA levels of gene set assigned to specified REACTOME Pathway.

Pathways highlighted in **Green** are significantly altered in both cell types; lipid-metabolism associated pathways (only altered in macrophages are highlighted in **Red**).

**357** (MDM) and **332** (monocytes) ranked pathways were returned with P values < 0.05.

**STable. 4.** SYBR qPCR and mouse genotyping primer sequences

| **Gene** | **Forward primer (5'→3')** | **Reverse primer (5'→3')** |
|---|---|---|
| *Abca1* | GGCCAGTCTGTGTAACGGAT | TGCATCGAGCTTCTTCCTCG |
| *Abcg1* | AGGTCTCAGCCTTCTAAAGTTCCTC | TCTCTCGAAGTGAATGAAATTTATCG |
| *Abdh5* | GTGTCCCCTGCACTTACAAGA | GGAGGACAAGTGGCGTCTTA |
| *Acat1* | ATTTGCTGACGCTGCTGTAGA | AAGGCTTCATTTACTTCCCACATTG |
| *β-actin* | GGGACCTGACAGACTACCTCATG | GTCACGCACGATTTCCCTCTCAGC |
| *Cd163* | CAGCGTAGTCTGCTCACGAT | AGTTGCTCCTGGCTGGTATG |
| *Cd68* | AAGGGGGCTCTTGGGAACTA | CGAAGGGATCGTCATAGCCC |
| *Cd36* | ATGGGCTGTGATCGGAACTG | GTCTTCCCAATAAGCATGTCTCC |
| *Ces1* | AGGGAGTTCTCGACGCAATG | ATGTAGGTGGGAGCTCCAGT |
| *Cxcl16* | GAGCGCAAAGAGTGTGGAACT | TGTGGACAAGGACCTGAAAAGT |
| *Hadhb* | ACCTGCGTTCATCAAACCCT | CAGAAGCGCCATCAGTCAGG |
| *Lpl* | TTGCAGAGAGAGGACTCGGA | GTTGCACCTGTATGCCTTGC |
| *Lxrα* | TCAGCATCTTCTCTGCAGACCGG | TCATTAGCATCCGTGGGAACA |
| *Marco* | GACAAGCCCTTCTTCTCGCT | CCAAGCCCTCTTTAAGCCCC |
| *Msr1* | CGCACGTGGAACAGGAAGTA | TCTGTGAGTGTTCCCAGTCC |
| *Nceh1* | AGAAGACCCTGCCACAATGA | CTGTGACTACGGGACACGATT |
| *Olr1* | AGATAGACACCCTCACCTTGAA | GTCACGCACGATTTCCCTCTCAGC |
| *Pltp* | CCATGTTGCCCGAAGTGGAG | CCCCGGAGTGTCAACTTAGC |
| *Pparγ* | GCCCTGGCAAAGCATTTGTA | TTTCCTGTCAAGATCGCCCT |
| *Ptgs1* | GTTCCGAGCCCAGTTCCAATA | AGCTGTACTCTTGTGAGCCC |
| *Scarb1* | GATGGAGAGCAAGCCTGTGA | GACTATGTGCAGGGGTGCTC |
| *Stab1* | ACGCTGACTGCCTCAATACC | AAGCATCAGTGTGGCAAGT |
| *Lyz2Cre* | Common:<br>CTTGGGCTGCCAGAATTTCTC | Mutant:<br>CCCAGAAATGCCAGATTACG<br><br>Wild type:<br>TTACAGTCGGCCAGGCTGAC |
| *Rosa26.Trib1* | Common:<br>GTGATCTGCAACTCCAGTCTTTC<br>TAG | Mutant:<br>CCTTCTTGACGAGTTCTTCTGAGG<br><br>Wild type:<br>CGCGACACTGTAATTTCATACTGTAG |
| *Trib1 fl/fl* | Common:<br>ACCTTGATCTGCAGTCCTAGG | Floxed:<br>AAGTTCACATTTGAACTGATGGC<br><br>Wild type:<br>AGCTGGTTTCAGGGGAAGAC |

**STable. 5. Linoleic (LiA), Oleic (OA) and Lauric Acid (LA) in vitro Polarised Human Monocyte Derived Macrophages[A] Recapitulate the *Olr1*$^{High}$/*Lpl*$^{High}$/*Scarb1*$^{Low}$/*CD36*$^{WT}$ RNA Profile *of Trib1*$^{mTG}$ BMDM.**

| | ***OLR1*** | | ***LPL*** | | ***SCARB1*** | |
|---|---|---|---|---|---|---|
| Stimulus | Fold Change | p | Fold Change | p | Fold Change | p |
| **LiA** | **6.36** | **6.60E-09** | **2.86** | **2.65E-05** | **-1.99** | **7.68E-07** |
| *SA* | 6.17 | 1.08E-08 | 3.19 | 4.00E-06 | -1.85 | 8.55E-06 |
| **OA** | **5.50** | **5.70E-12** | **2.65** | **3.87E-07** | **-1.38** | **1.66E-03** |
| **LA** | **5.20** | **1.73E-07** | **4.44** | **6,37E-09** | **-1.33** | **3.32E-02** |
| *PA* | 4.76 | 1.78E-10 | 5.62 | 7.26E-17 | -1.75 | 1.31E-07 |
| *TNF_PGE2* | 4.42 | 3.88E-04 | -2.10 | 2.58E-02 | -1.52 | 2.33E-02 |
| TNF_P3C | 3.86 | 1.21E-03 | -1.38 | 3.30E-01 | -1.24 | 2.34E-01 |
| upLPS | 3.42 | 3.40E-06 | 1.82 | 4.15E-03 | -1.12 | 3.14E-01 |
| *TPP* | 2.66 | 9.76E-05 | -2.80 | 5.33E-07 | -1.38 | 3.53E-03 |
| *TPP_IFNb* | 2.37 | 3.68E-02 | -3.28 | 4.31E-04 | -1.62 | 8.93E-03 |
| *TNF* | 2.30 | 6.50E-03 | 1.86 | 1.12E-02 | -1.68 | 1.61E-04 |
| *sLPS* | 1.86 | 2.10E-02 | -1.36 | 1.60E-01 | -1.27 | 4.43E-02 |
| sLPS_IFNg | 1.80 | 1.54E-01 | -2.00 | 3.71E-02 | -1.62 | 8.86E-03 |
| IFNg | 1.73 | 5.39E-02 | 1.33 | 2.10E-01 | -1.51 | 1.17E-03 |
| IFNg_TNF | 1.49 | 3.30E-01 | 2.14 | 2.20E-02 | -1.92 | 4.33E-04 |
| sLPS_IC | 1.46 | 3.57E-01 | -1.49 | 2.27E-01 | -1.38 | 8.14E-02 |
| HDL | 1.33 | 4.31E-01 | -1.17 | 5.81E-01 | -1.02 | 8.94E-01 |
| IFNb | 1.25 | 5.91E-01 | 1.45 | 2.58E-01 | -1.07 | 7.25E-01 |
| TPP_IFNb_IFNg | -1.01 | 9.81E-01 | -3.52 | 1.95E-04 | -1.91 | 4.74E-04 |
| IL10 | -1.04 | 9.31E-01 | 2.74 | 2.59E-03 | 1.09 | 6.24E-01 |
| P3C_PGE2 | -1.18 | 6.90E-01 | -1.84 | 6.58E-02 | -1.33 | 1.14E-01 |
| P3C | -1.33 | 4.93E-01 | -1.03 | 9.23E-01 | 1.00 | 9.91E-01 |
| PGE2 | -1.37 | 4.46E-01 | -1.59 | 1.63E-01 | 1.08 | 6.75E-01 |
| upLPS_IC | -1.44 | 3.76E-01 | 1.03 | 9.24E-01 | 1.23 | 2.53E-01 |
| IL13 | -1.97 | 9.89E-02 | 3.52 | 1.92E-04 | -1.37 | 8.55E-02 |
| GC | -2.28 | 4.59E-02 | -3.33 | 3.62E-04 | 1.38 | 8.13E-02 |
| IL4_upLPS | -2.37 | 4.74E-03 | 1.61 | 5.12E-02 | 1.36 | 2.14E-02 |
| IL4 | -3.04 | 1.49E-06 | 2.05 | 9.04E-05 | 1.03 | 7.69E-01 |
| TRIB1 high MDM | 1.86 | 4.76E-16 | 1.43 | 6.32E-11 | -1.18 | 2.00E-08 |

[A]Data extracted from Xue *et al.*

*Italics indicate the eleven conditions that produce reciprocal changes in OLR1 and SCARB1 RNA.*

***Bold** indicates the three conditions that recapitulate the Olr1*$^{High}$/*Scarb1*$^{Low}$/*Lpl*$^{High}$ profile of *Trib1*$^{mTg}$ BMDMs.
